# Spliceosome-Associated MicroRNAs Identified in Breast Cancer Cells Act on Nuclear Targets and Are Potential Indicators for Tumorigenicity

**DOI:** 10.1101/2020.07.19.211292

**Authors:** Shelly Mahlab-Aviv, Keren Zohar, Yael Cohen, Ayelet R. Peretz, Tsiona Eliyahu, Michal Linial, Ruth Sperling

**Affiliations:** School of Computer Science and Engineering, The Hebrew University of Jerusalem, Jerusalem, Israel; Department of Biological Chemistry, The Hebrew University of Jerusalem, Jerusalem, Israel; Department of Genetics, The Hebrew University of Jerusalem, Jerusalem, Israel

**Keywords:** RNA-seq, AntimiR, OncomiR, carcinogenesis, ncRNA, Metastasis

## Abstract

MicroRNAs (miRNAs) act as negative regulators of gene expression in the cytoplasm. Previous studies identified miRNAs associated with the spliceosome. Here we study three breast-derived cell-lines with increased tumorigenicity (from MCF-10A to MCF-7 and MDA-MB-231) and compared their miRNA sequences at the spliceosome fraction (SF). We report that the SF-miRNAs expression, identity, and pre-miRNA segmental composition vary across these cell-lines. The expression of the majority of the abundant SF-miRNAs (e.g. miR-100, miR-30a, and let-7 members) shows an opposite trend in view of the literature and breast cancer large cohorts. The results suggest that SF-miRNAs act in the nucleus on alternative targets than in the cytoplasm. One such miRNA is miR-7704 whose genomic position overlaps HAGLR, a cancer-related lncRNA. We found an inverse expression of miR-7704 and HAGLR in the tested cell lines. Moreover, inhibition of miR-7704 caused an increase in HAGLR expression. Furthermore, increasing miR-7704 levels attenuated the MDA-MB-231 cell-division rate. While miR-7704 acts as oncomiR in breast cancer patients, it has a tumor-suppressing function in SF, with HAGLR being its nuclear target. Manipulating miR-7704 levels is a potential lead for altering tumorigenicity. Altogether, we report on the potential of manipulating SF-miRNAs as an unexplored route for breast cancer therapeutics.

## 1. Introduction

MicroRNA (miRNAs) are small, ∼22 nt long, non-coding RNA molecules implicated in defining cell identity and regulating signaling pathways. Variations in miRNA expression have been linked with numerous human diseases including cancer [1, 2]. The main studied role of miRNAs in mammals is in inhibiting translation and negatively controlling gene expression. This is mediated by base-pairing of miRNAs primarily to the 3’-UTR of the target mRNA transcripts in the cytoplasm [3-7]. In humans, a large fraction of miRNA genes is located in gene introns. The canonical biogenesis of intronic miRNAs from Pol II transcripts involves two main steps: The first step occurs in the nucleus by the microprocessor, whose key proteins are DGCR8 and Drosha. DGCR8 binds the RNA molecule, while Drosha, an RNase III type enzyme, cleaves the primary (pri) miRNA transcript into a precursor (pre) miRNA stem-loop molecule of 70-80 bases [8-12]. The second step occurs after the export of the pre-miRNA to the cytoplasm [13] when it is cleaved by Dicer yielding a mature miRNA, which is loaded on the RNA Induced Silencing Complex (RISC) [3, 4, 14]. The RISC-bound miRNAs act in targeting mRNAs in the cytoplasm [7]. According to miRBase [15] over 1900 miRNA genes are found in human genome, yielding ∼2600 mature miRNAs. It was estimated that miRNAs control ∼60% of all human genes [3]. Alternation in the quantity of miRNAs expression has the potential to perturb cellular pathways leading to human diseases such as neurodegenerative diseases, chronic inflammation, metabolic disorders, and cancer onset and progression [1, 2, 15, 16].

The finding of mature miRNAs in the nucleus [17-21], suggested nuclear functions for miRNA in addition to the classical cytoplasmic ones [22, 23]. Active process of shuttling of miRNAs from the cytoplasm to the nucleus were demonstrated [24, 25]. The function of miRNAs in the nucleus are not yet well understood. However, the involvement of miRNA in a number of processes, such as the regulation of non-coding RNAs [26-30], transcriptional silencing [31, 32], activation [33-36], and inhibition [32] were reported. Furthermore, analysis of miRNA-mRNA-AGO interactions, revealed substantial AGO-miRNA mapping to intronic sequences [37].

A large fraction of miRNA genes is located in introns of coding genes, while many are expressed from their own Pol II promoters [6, 38, 39]. For most intronic miRNAs, the mRNA and such miRNA can be expressed from the same primary transcript. However, when the pri-miRNA is located in an exon or overlaps a splice site, at any specific time only the mRNA or the miRNA can be generated from the single transcript. In the case of alternative splicing (AS), the expression of both the mRNA and the miRNA from the same transcript is possible [39]. Furthermore, several studies showed links between splicing and miRNA processing [25, 40-44].

Splicing and AS play a major role in the regulation of gene expression in mammals [45]. Furthermore, changes in AS occur in many human diseases including cancer [46-48]. Splicing occurs in the cell nucleus within a huge (21 MDa) and highly dynamic machine known as the supraspliceosome [49, 50]. It is composed of four native spliceosomes, which are connected by the pre-mRNA [51, 52]. The entire repertoire of nuclear pre-mRNAs, independent of their length and number of introns, is individually assembled in supraspliceosomes [49, 50]. The supraspliceosome offers coordination and regulation of pre-mRNA processing events. Thus, it is involved in all nuclear processing activities of pre-mRNAs [49, 50]. Furthermore, miRNAs were found within the endogenous spliceosome [44, 53-55], where a cross-talk between pre-mRNA splicing and miRNA processing was demonstrated [44, 55].

A recent study on the composition of small ncRNA (<200 nt) within the supraspliceosomal fraction (SF) from HeLa cells, identified about 200 miRNA sequences [54], several that are exclusively expressed in the SF. Many of these sequences are not harbored in introns. These findings indicate that the presence of miRNA sequences in the endogenous spliceosome is not only due to biogenesis, but is suggesting novel functions for miRNAs in the endogenous spliceosome. While about two thirds of these miRNA sequences were identified as mature miRNAs, the rest represent segments derived from pre-miRNAs. One of these exclusive SF-miRNA is miR-7704 that was shown to be a negative regulator of HAGLR (HOXD Antisense Growth-Associated Long Non-Coding RNA) expression in HeLa cells [54].

In this study, we focused on the expression of SF-miRNAs in human breast cancer cell-lines. We analyze SF-miRNA in malignant cell-lines and a non-tumorigenic cell-line and determine the SF-miRNAs composition and their unique properties. We investigated miR-7704 and show that it negatively regulates the expression of lincRNA HAGLR, which plays a role in the development and progression of multiple cancers [56]. The potential role of SF-miRNAs in regulating breast cancer progression is discussed.

## 2. Results

### 2.1. Isolation of Spliceosomal RNA from Breast Cell-Lines

The identification of hundreds of miRNA sequences in SF from HeLa cells, with several of them known to play a role in cancer, prompted us to search for spliceosomal miRNAs in breast cancer cells. To this end we have chosen the breast cell-line MCF-7, an estrogen-dependent ductal carcinoma, and the MDA-MB-231 which is a highly invasive and metastatic estrogen-independent adenocarcinoma. We also tested MCF-10A which is a non-tumorigenic mammary epithelial cell-line, as a model for normal breast cell function. We prepared nuclear supernatants enriched with supraspliceosomes under native salt conditions from the three cell-lines, and fractionated each on glycerol gradients as previously described [51], (see Materials and Methods). Notably, the isolation protocol preserves the higher order splicing complexes as shown by electron microscopy [51, 57]. The supraspliceosomes sediment at 200S with the splicing factors SR proteins, hnRNP G, and REF/Aly, as well as the cap binding protein CBP80. We used these components to locate the position of the supraspliceosome in these gradients [58-60]. Figure 1 presents the results of Western blot analyses across the glycerol gradient performed on samples from each of the breast cell-lines, using antibodies directed against the above genuine supraspliceosome components.

**Figure 1.**
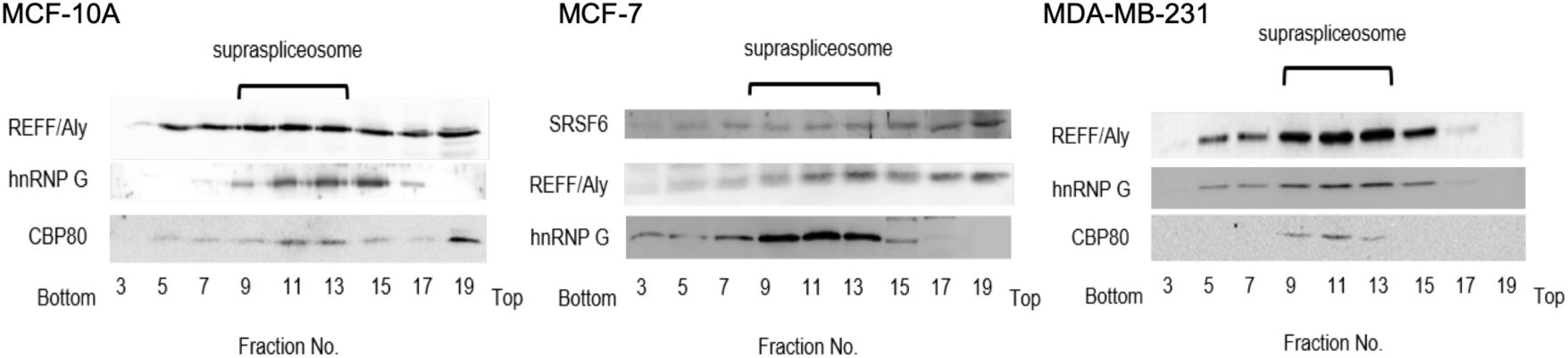
Splicing factors mark the supraspliceosome fraction (SF) in breast cancer cells. Western blot analysis of the distribution across the glycerol gradient of splicing factors. Nuclear supernatants enriched for SF were prepared from MCF-10A (**A**), MCF-7 (**B**), and MDA-MB-231 (**C**), and were fractionated in 10-45% glycerol gradients. Tobacco mosaic virus (TMV) was used as a size marker for the sedimentation. Aliquots from odd gradient fractions were analyzed by Western Blot using anti-hnPNP G, anti-SR, anti-REF/Aly and anti CBP80 antibodies. Supraspliceosomes peak in fractions 9-13 (200S). The distribution of hnRNP G (42 kDa), SRSF6 (SRp55, 55kDa), REF/Aly (27 kDa), and CBP80 (80 kDa) across the gradient is shown.

Next, we extracted small RNA (<200 nt) from the spliceosomal fraction (SF, fractions 9-12, Figure 1) of each of the breast cell-lines and used the RNA in the SF for constructing a barcoded library of small RNA for further sequencing, as previously described [44, 54]. We created three different libraries for biological triplicates from each cell-line. Statistical summary of the RNA-seq libraries is shown in Supplementary Table S1. Alignment of these SF sequences to the human transcriptome revealed a complex collection of RNA species, including pre-miRNAs, small nucleolar RNAs (SNORDs) [61], intronic sequences and more. In this study, we only consider reads that are aligned to the hairpin precursor miRNA of the primary miRNA as determined by miRbase [15]. These sequences are referred to as SF-miRNAs.

### 2.2. Changes in Expression of SF-miRNA Sequences in Breast Cancer Cells

Sequencing and alignment to all transcriptome and miRNA collection revealed a large complexity of the miRNA sequences with 397, 195 and 246 different entities for MCF-10A, MCF-7 and MDA-MB-231 cells, respectively (≥2 reads per cell-line). Note that the amounts of SF-miRNAs aligned sequence reads range over 2-3 order of magnitude (Supplementary Table S2).

To increase reliability, we restricted the analysis to the subset of higher confidence sequences (read length ≥17 nt, ≥10 total reads from each cell-line). Figure 2 shows a heatmap of the quantities of the different SF-miRNA from each of the experimental RNA-seq libraries (3 cell-lines in triplicates; see Materials and Methods). We further tested the correlation of the miRNAs identified for each of the 9 sets of miRNA profiles (Supplemental Figure S1). Note that the triplicates of MCF-10A show the stronger correlation, and the correlations of the miRNA profile for MCF-7 and MDA-MB-231 are less distinctive.

**Figure 2.**
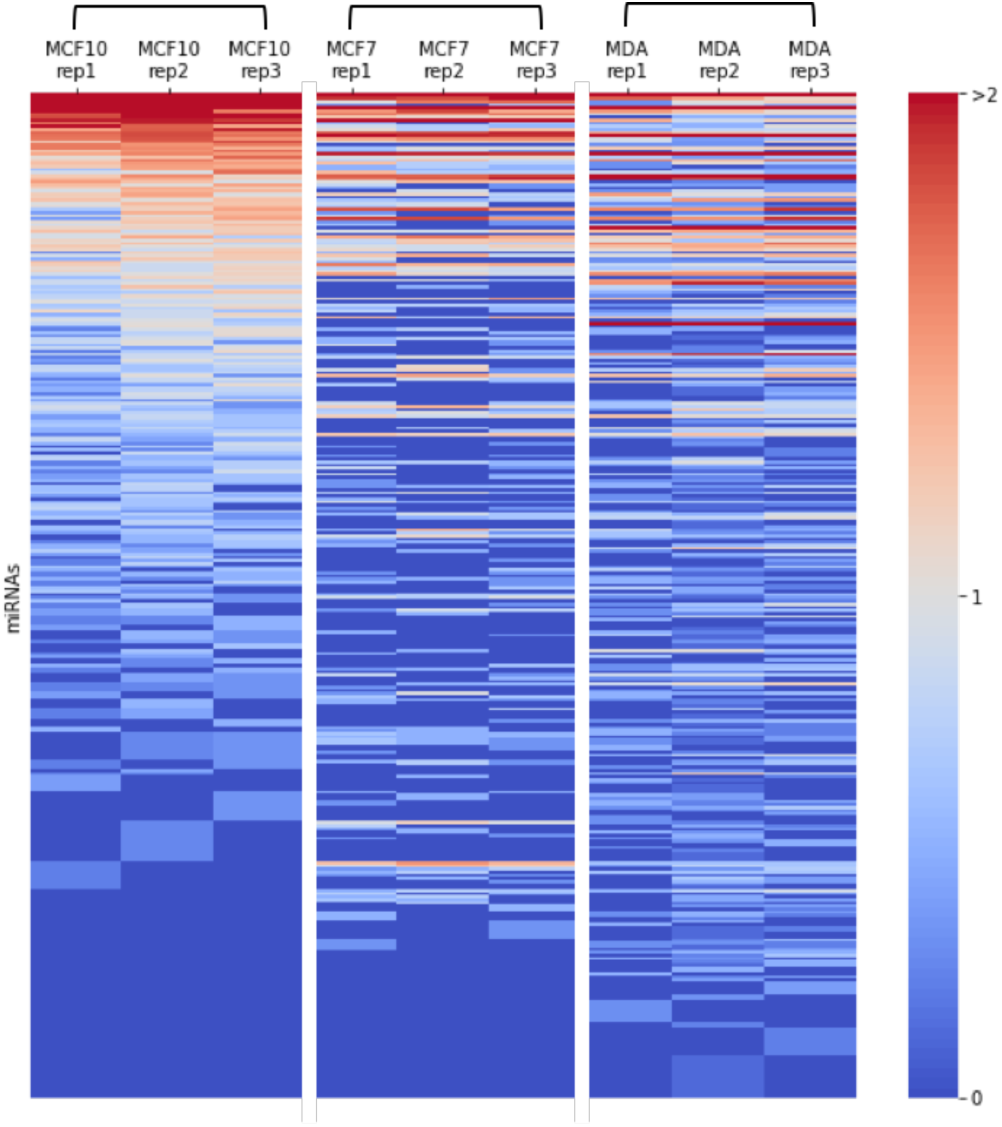
Heatmap of miRNA expression in breast cancer cells. Expression of SF-miRNAs in triplicates from each of the breast cancer cell-lines: MCF-10A, MCF-7, and MDA-MB-231, are presented. Color code represents the amount of reads in logarithmic scale (log10). All miRNAs that are analyzed have ≥10 reads in any specific cell type, a minimal read length of ≥17. Data source is in Supplemental Table S2.

Based on the strict thresholds on reliable reads, (with a threshold of 10 total reads per cell-line) we identified 155, 56 and 102 SF-miRNAs in MCF-10A, MCF-7 and MDA-MB-231 cells, respectively. The unified list includes reliable finding for 191 SF-miRNAs (Supplementary Table S2). Figure 3A presents a Venn diagram of the overlap of SF-miRNAs found in the breast cancer cell-lines. We find that 45 miRNA sequences are expressed in all three cell-lines. The highest expressing miRNAs include miR-6087, miR-21, miR-20a and let-7g (Supplemental Table S2). When testing the miRNAs that are shared in both MCF-10A and MCF-7, additional 3 miRNAs were found: miR-3652, miR-5047, and miR-200c. Additional 24 SF-miRNAs are shared between MCF-10A and MDA-MB-231 (e.g. miR-100, miR-222, miR-221 and miR-30a). For the two cancerous cell-lines (MCF-7 and MDA-MB-231) additional 5 SF-miRNAs were found (e.g. miR-492, miR-200b). Figure 3A further illustrates that many of the miRNAs are exclusively expressed in a specific cell-line. A total of 83, 3, and 28 are expressed solely in MCF-10A, MCF-7 and MDA-MB-231, respectively.

**Figure 3.**
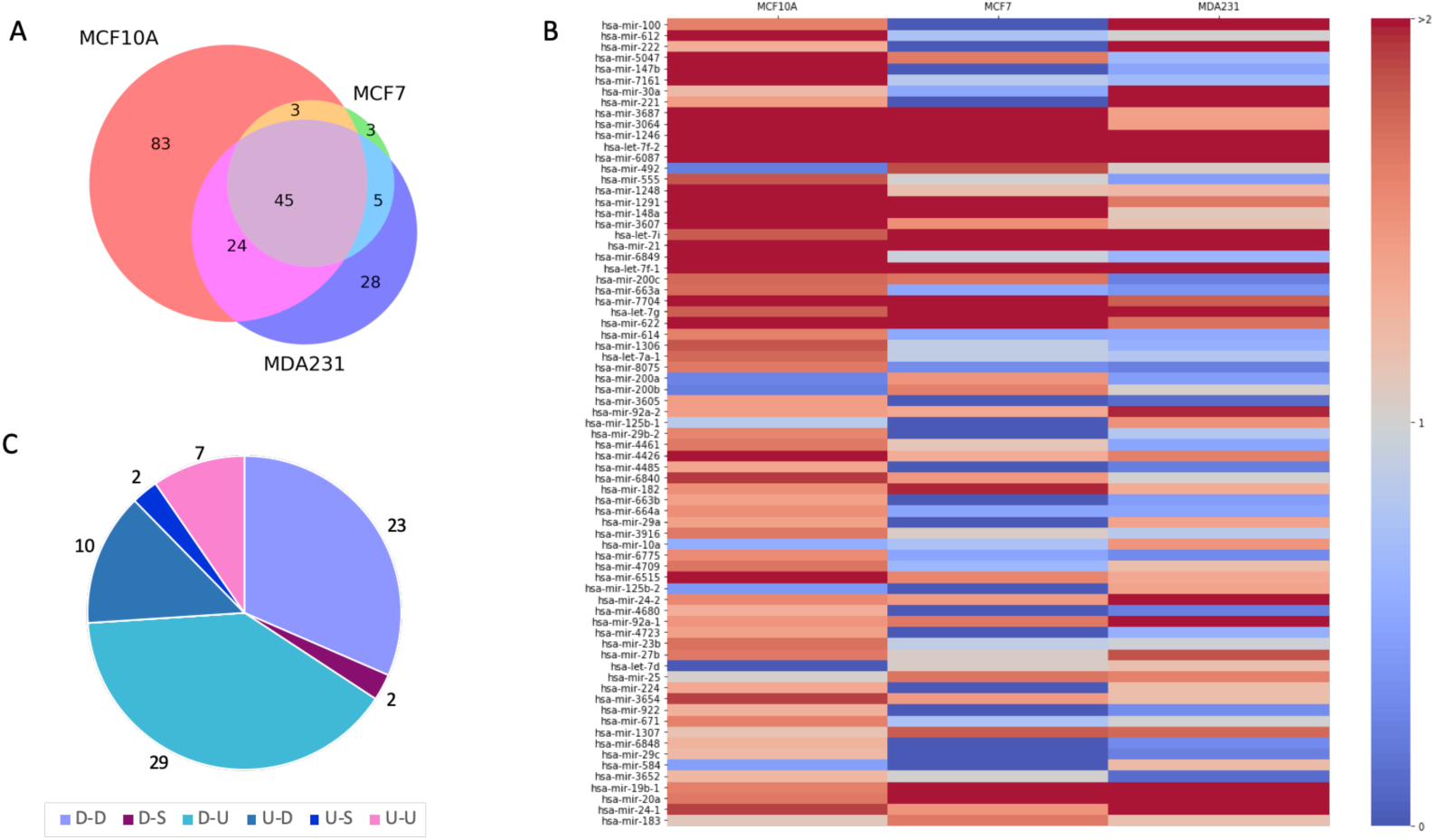
Changes in expression of SF-miRNAs among three breast cell-lines. **(A)** Venn diagram showing the partition of different SF-miRNAs among three breast cancer cell-lines. Source data from Supplementary Table S2. **(B)** Following normalization of reads and testing statistics for differential expression across the three cell-lines, a significant list of 73 SF-miRNAs is shown, colored by the amounts of normalized reads (log 10). All listed 73 SF-miRNAs met the statistical adjusted p-value of <0.05 and are ranked by the statistical significance. **(C)** Partition of the 73 SF-miRNA according to the 9 possible paired expression trends, composed of up (U), down (D) and S (same). Paired expression trend is the trend for MCF-7 versus MCF-10A and then MDA-MB-231 versus MCF-7. The dominant observed trend is marked as D-U followed by D-D. Note that only 6 out of 9 possible paired trends were observed. Data source is in Supplemental Table S3. Expression of SF-miRNAs in triplicates from each of the breast cancer cell-lines: MCF-10A, MCF-7, and MDA-MB-231, are presented. Color code represents the amount of reads in logarithmic scale (log10). All miRNAs that are analyzed have ≥10 reads in any specific cell type, a minimal read length of ≥17. Data source is in Supplemental Table S2.

For testing the trend in expression of each of SF-miRNAs, we analyzed the differential expression profile by the statistical significance of the expression results following normalization (DEseq2, Supplemental Table S3, see Materials and Methods). Figure 3B shows the results by a heatmap with colors indicating the level of expression (in log scale). The miRNAs are sorted according to the differential expression significant findings. Specifically, the most significant differential expressed SF-miRNA across all three cell-lines is miR-100 (adjusted p-value = 4.65E-65). Only significant miRNAs with adjusted p-value <0.05 are listed. All together there are 73 such miRNAs (Supplemental Table S3).

Figure 3C presents a partition of expression trends of the 73 listed SF-miRNAs that show significant alterations in the expression among the different cells (Figure 3B). Specifically, differences in expression of miRNAs are tested in view of the elevated cancer characteristics of the cells. We compared MCF-7 relative to MCF-10 and further MDA-MB-231 relative to MCF-7. We transformed expression levels to discrete trends marked as Up (U), Down (D) and Same (S). All together there are 9 possible such paired trends. For example, U-D indicates an increase in MCF-7 relative to MCF-10A and a decrease in MDA-MB-231 relative to MCF-7. A trend marked as Same indicates expression that is within ±10% change in expression. The partition of the observed trends is shown in Figure 3C. The dominant trends (54 miRNAs, 74%) are a decrease in expression when comparing MCF-7 to MCF-10A. Among them, the majority (29 of 54 miRNAs) show a trend of D-U. The rest of the SF-miRNAs (26%, 19 miRNAs) marked as increased in the MCF7 relative to MCF-10A (Figure 3C).

Table 1 presents the top 25 SF-miRNAs ranked by their expression levels in the three cell-lines (see also Supplementary Table S3). The total expression levels of the different miRNAs for each of the cell-lines is shown along with their associated expression trends. Table 1 also includes the genomic position of the identified SF-miRNAs. Many of these miRNA-aligned sequences are intergenic, therefore their appearance is not a mere reflection of intronic pre-miRNAs processing of transcribed host genes. For some miRNAs (miR-1291, miR-1248, miR-3607) their genomic location overlaps SNORA/Ds. For 8 of the listed miRNAs, the position of the sequences is in the introns of coding genes. The majority of the other miRNAs are associated with lncRNAs, including transcripts that accounts for clusters of neighboring miRNAs (Table 1). Several miRNAs that belong to the same genomic cluster were detected as SF-miRNAs (e.g., miR-5047 and miR-3064, Table 1). Among these differential miRNAs some are known as oncogenic (e.g., miR-21, miR-222, mir-221 and miR-1246) and others as tumor suppressors (e.g., let-7, miR-30a and miR-100). We conclude that many of the identified SF-miRNAs are found at the vicinity of ncRNA and in regions of miRNA within local genomic clusters.

**Table 1.** Highly expressed 25 SF-miRNAs.

| miRNA<sup>a</sup> | Mean MCF10 | Mean MCF7 | Mean MDA | Exp. Trend<sup>d</sup> | Genomic annotation | Cluster neighbor<sup>c</sup> |
| --- | --- | --- | --- | --- | --- | --- |
| hsa-mir-6087* | 5847.1 | 1124.3 | 705.3 | D-D | Intergenic |  |
| hsa-mir-21 | 362.6 | 1173.3 | 1155.5 | U-S | Exon, 3'UTR |  |
| hsa-mir-1246 | 484.6 | 83.8 | 32.0 | D-D | Intron, LncRNA |  |
| hsa-mir-3687* | 460.6 | 34.9 | 8.2 | D-D | Ribosomal RNA |  |
| hsa-mir-100 | 13.1 | 0.0 | 483.4 | D-U | Intron, LncRNA | hsa-mir-526; hsa-let-7a-2 |
| hsa-mir-612 | 450.5 | 1.6 | 3.0 | D-U | LncRNA |  |
| hsa-mir-7704 | 296.4 | 41.8 | 20.2 | D-D | Exon, LncRNA |  |
| hsa-let-7f-1 | 54.0 | 90.6 | 184.7 | U-U | Intergenic | **hsa-let-7a-1**; hsa-let-7d |
| hsa-let-7f-2 | 32.1 | 86.0 | 173.2 | U-U | Intergenic | **hsa-mir-98** |
| hsa-let-7g | 20.4 | 84.8 | 170.6 | U-U | Intron, Coding |  |
| hsa-mir-3064 | 196.6 | 46.3 | 8.8 | D-D | Mirtron, Coding | **hsa-mir-5047** |
| hsa-mir-1291** | 88.0 | 147.6 | 14.9 | U-D | Intron, Coding |  |
| hsa-let-7i | 21.2 | 62.3 | 140.3 | U-U | Mirtron, LincRNA |  |
| hsa-mir-5047 | 163.8 | 14.5 | 1.2 | D-D | Intron, Coding | **hsa-mir-3064** |
| hsa-mir-622 | 54.6 | 74.2 | 16.8 | U-D | Intergenic |  |
| hsa-mir-19b-1 | 14.0 | 60.1 | 50.6 | U-D | LncRNA | hsa-mir-17; hsa-mir-18a; hsa-mir-19a; **hsa-mir-20a**; **hsa-mir-92a-1** |
| hsa-mir-20a | 15.1 | 53.2 | 51.0 | U-S | LncRNA | hsa-mir-17; hsa-mir-18a; hsa-mir-19a; **hsa-mir-19b-1**; **hsa-mir-92a-1** |
| hsa-mir-222 | 6.5 | 0.0 | 105.5 | D-U | Intron, LncRNA | **hsa-mir-221** |
| hsa-mir-30a | 5.2 | 0.8 | 91.1 | D-U | Intron, LincRNA |  |
| hsa-mir-148a | 47.4 | 32.4 | 4.0 | D-D | Intergenic |  |
| hsa-mir-7161 | 74.7 | 1.9 | 1.2 | D-D | Intron, Coding |  |
| hsa-mir-24-1 | 27.1 | 10.2 | 40.3 | D-U | Intron, Coding | **hsa-mir-23b**; **hsa-mir-27b**; hsa-mir-3074 |
| hsa-mir-1248** | 66.8 | 4.5 | 5.3 | D-U | Intron, Coding |  |
| hsa-mir-3607** | 57.0 | 10.8 | 4.3 | D-D | Intron, Coding |  |
| hsa-mir-221 | 8.5 | 0.0 | 59.7 | D-U | Intron, LncRNA | **hsa-mir-222** |

<sup>a</sup>miRNAs marked by \* were removed from the recent version of miRbase (ver22.0); miRNAs marked by \*\* overlaps with SNORA/D.
<sup>b</sup>Expression trend: U, D and S depicts up, down and same, respectively. The paired trend represents the relative level of miRNA expression in MCF-7 relative to MCF-10 and MDA-MB-231 relative to MCF-7. Same indicates stable expression within $\pm 10\%$ .
<sup>c</sup>Cluster neighbors are miRNAs within a genomic distance <10k, as defined by miRbase. Marked in bold are cluster neighbor miRNAs which are found among SF-miRNAs (supported by >10 reads).

### 2.3. Changes in the Segmental Regions of SF-miRNA in Breast Cancer Cell-Lines

To further characterize the properties of the SF-miRNAs we assessed the segmental composition of each identified miRNA. We observed that the SF-miRNA sequences are not only confined to mature miRNAs. Therefore, other sequences from the pre-miRNA may act on nuclear targets by base-pair complementarity. We classified the SF-miRNA sequences according to the segmental regions to which they align (Figure 4A). These regions are based on the complete hairpin precursor miRNA (HP-miRNA) sequence (as determined by miRBase), mature miRNA (either derived from the 5p or the 3p); undefined complement (i.e. a complementary sequence of mature miRNA where there is no experimental evidence in miRBase), and miRNA extension (tails of <50 nt of genomic extended sequences from the mature miRNA / complementary sequence). In addition, reads that do not reside within any of these predefined regions, but overlap two or more regions, are categorized as overlapping region.

**Figure 4.**
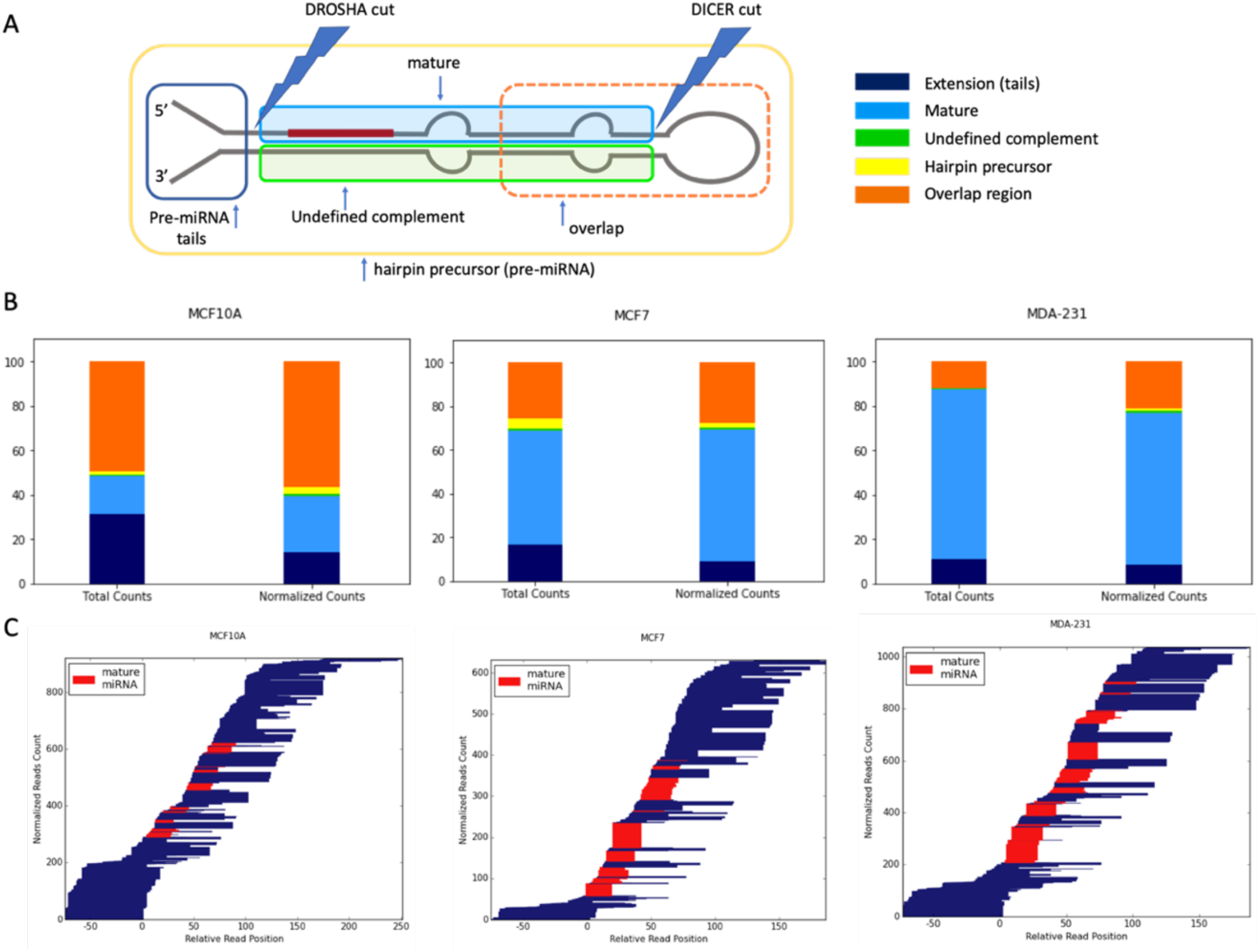
Partition of the SF-miRNAs to segmental pre-miRNA regions. **(A)** A schematic view of a miRNA prototype, where the different regions of the pre-miRNA are listed according to their positions on the pre-miRNA major processing sites. An example of overlap regions, defined as reads that cross known segmental boarder, is indicated. **(B)** The relative counts of reads to each of the pre-miRNA regions. Total counts of reads (left) and normalized counts (right, the relative abundance of reads aligned to a specific miRNA sum to 1) by each miRNA are shown for MCF-10A, MCF-7 and MDA-MB231. The sum of reads mapped to all the reads according to the cell-lines. (**C)** Blocks of reads from the same starting points reflect non-randomized cleavage sites. The 5’p and 3’p mature miRNAs are colored in red, and the width of the bars is proportional to the number of reads. The position and the alignment of the mature miRNAs is as annotated by miRBase. All miRNAs that are analyzed have an average minimal number of reads per cell-line (≥3), and a minimal read length of ≥17. Data source is in Supplemental Table S4.

Figures 4B show the distribution of reads among these disjoined categories in the three tested cell-lines (see also Supplemental Table S4). It can be seen that there is a major shift in the type of sequences between the non-malignant MCF-10A breast cells in which the mature miRNAs accounts for only ∼25% (length size ∼22 nt). In this cell-line ∼14% of the aligned reads represent extension sequences while the majority (∼57% of normalized counts) belong to overlapping regions. However, in the MCF-7 cells, the percentage of mature miRNA increases to ∼60% and reach over 68% of the sequences in MDA-MB-231 cells (Figures 4B and Supplemental Table S4). The percentage of overlap regions shows the opposite trend where the fraction in MCF-10A in maximal (57%) and lowest for MDA-MB-231 (21%). The fraction of full-length SF-pre-miRNA is rather low and decreased from 3% to 1% from MCF-10A to mDA-MB-231, respectively. The remaining miRNA aligned sequences belong to miRNA extension categories that decreased from 14% to 9% from MCF-10A to MDA-MB-231. Notably, the distribution along the pre-miRNA sequence is quite wide as demonstrated in Figures 4C. The scheme shows that the amounts, length and segmental profiles of the pre-miRNA genomic sequence varies substantially among the three cell-lines. In most SF-RNAs from MDA-MS-231 cells the mature miRNAs dominate. However, some display a more complex segmental composition. For example, the aligned reads for miR-151a, miR-181a-1, miR-146a and miR-98 were split between mature and overlap regions. Segmental regions of SF-miRNAs are listed in Supplemental Table S4.

### 2.4. Comparing the Differential Expression of SF-miRNAs in Breast Cell-lines with Respect to Cancer Literature

Figure 5A shows the log scale expression of the top SF-miRNAs in the cell-lines that specify non-malignant (MCF-10A), malignant (MCF-7) and highly metastatic cancerous cells (MDA-MB-231). Note that for a number of cases, orders of magnitude changes in expression level are recorded across the three tested cell-lines. Figure 5B shows the partition of expression for each miRNA among the three cell-lines. Two groups of SF-miRNAs are evident, those showing a directional decrease in abundance with increased malignancy, and the opposite trend of a decrease in abundance with malignancy.

**Figure 5.**
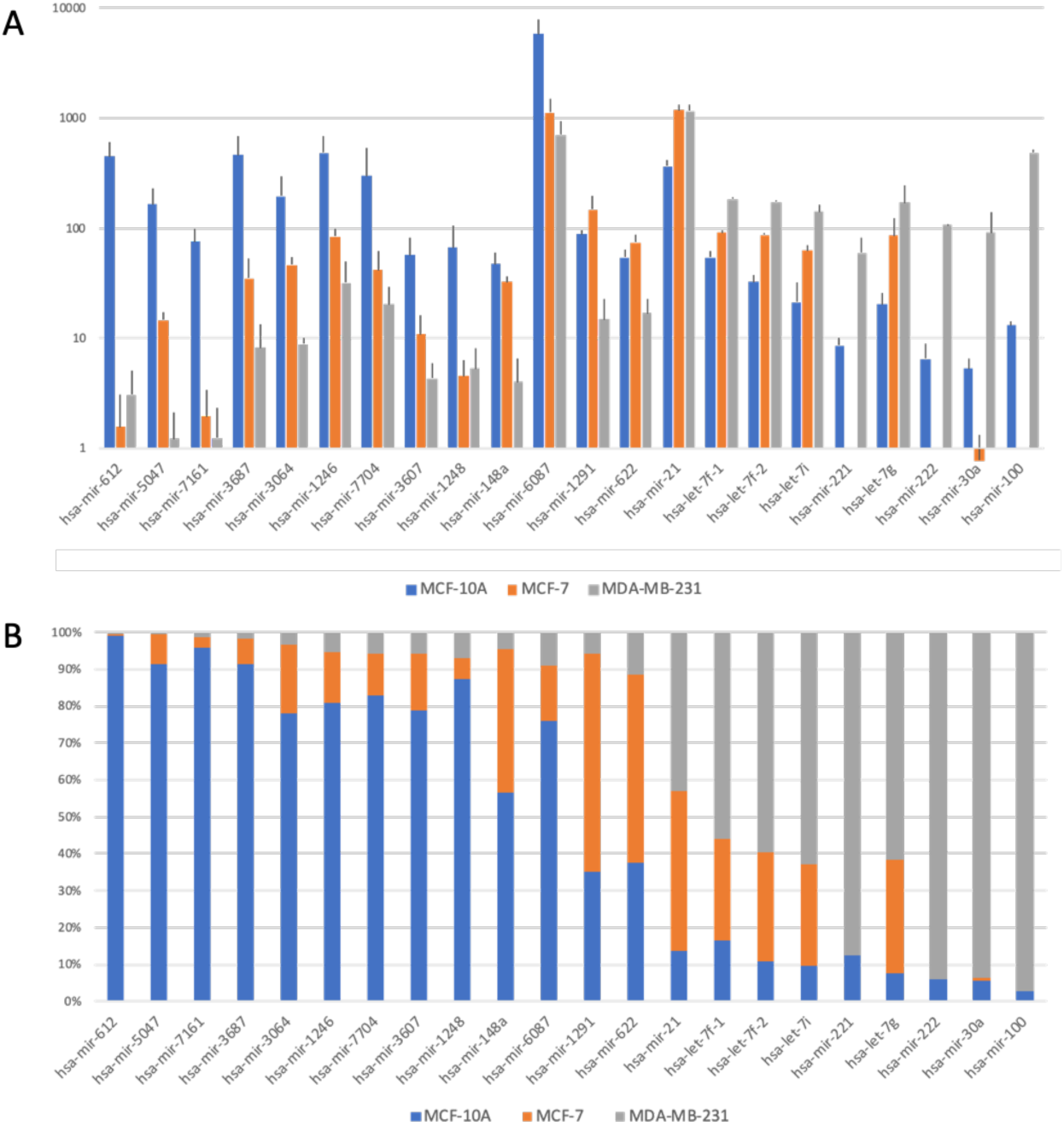
Changes in expression of top SF-miRNAs with breast cancer tumorigenicity. **(A)** Changes in the expression level of the top 22 SF-miRNAs in the three-breast cell-lines (log scale). **(B)** The partitions for each miRNA within the three cell-lines is shown. MCF-10A (blue), MCF-7 (orange) and MDA-MB-231 (grey). Data source is in Supplemental Table S3.

The larger differences are associated with the non-malignant (MCF-10A) and the metastatic cell-line (MDA-MB-231). Therefore, we compare the fold change and the trend in expression of SF-miRNAs between these two cell-lines (Figure 6). We limited the analysis to significant SF-miRNAs (Supplementary Table S3) with a joint expression level >50 normalized reads (total 22 miRNAs). Figure 6 also shows the consistency in the trend of expression with respect to the current literature-based knowledge as annotated by miRCancer [62]. miRCancer compiles publications on cancer and annotate each by the miRNA trend in cancerous versus healthy samples (marked Up / Down). While for several SF-miRNAs there is no evidence in miRCancer (Figure 6, stripped bar), others are consistent with the miRCancer report (e.g. miR-21, miR-148a, green). Importantly, for a number of miRNAs, an opposite trend is monitored for cytoplasmic miRNAs compared with the SF-miRNAs (Figure 6, red). The expression of miR-100, miR-30a, let-7g, let-7i, let-7f-1 and let-7f-2 is consistently suppressed in breast cancer samples, while the expression level of these miRNAs among the SF-miRNAs increases in MDA-MB-231 relative to MCF-10A. We propose that the observed increased SF-miRNAs expression in these apparent tumor suppressor genes (i.e. miR-100, miR-30a, let-7g, let-7i, let-7f-1 and let-7f-2) might serve as molecular indicators for cancerous severity in humans. These findings are consistent with the possibility that these SF-miRNAs are acting on nuclear targets which are likely different from the known cytoplasmic ones. For miRCancer reports on the selected SF-miRNA see Supplemental Table S5.

**Figure 6.**
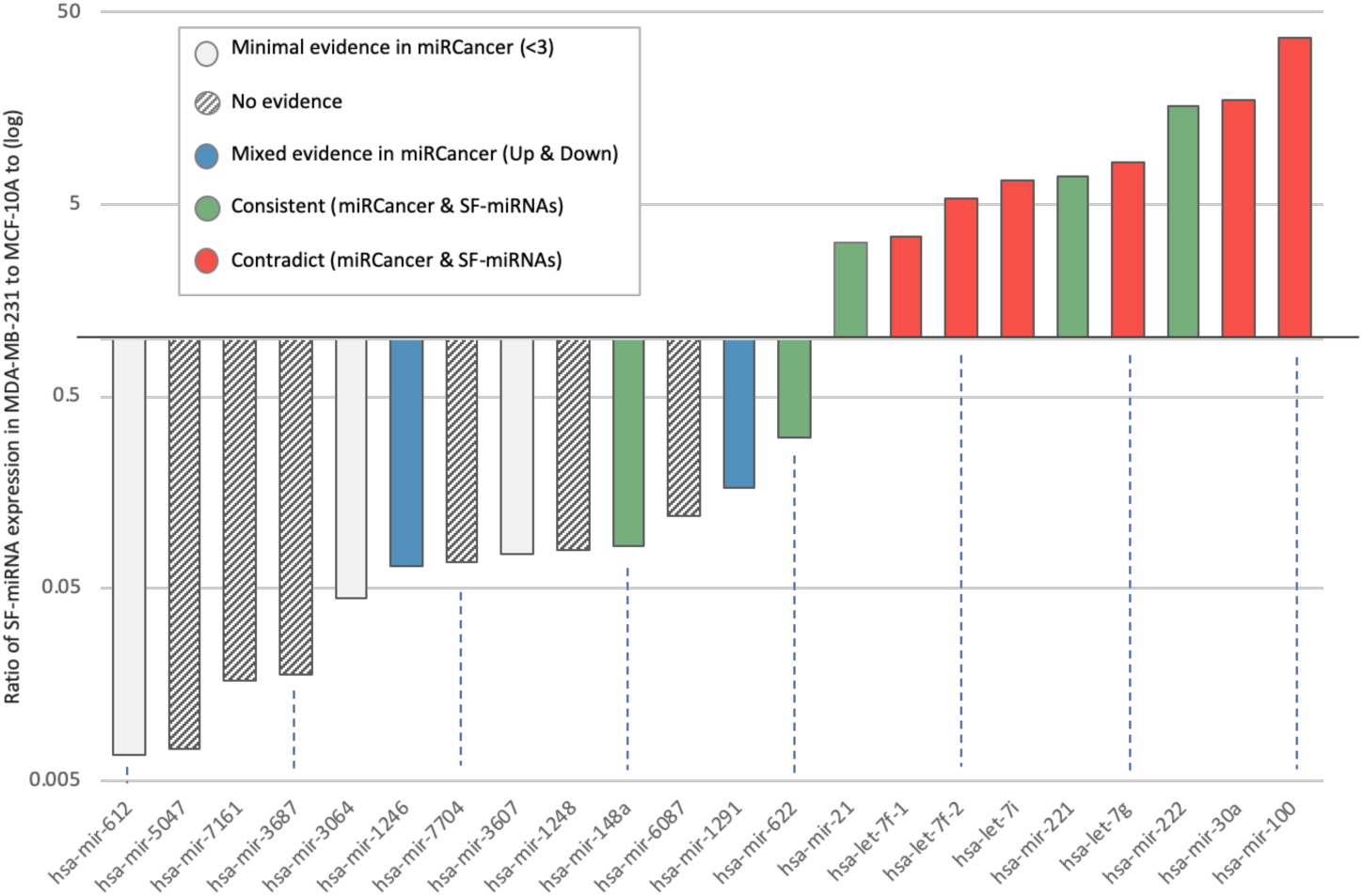
Expression level of SF-miRNAs in breast cancer cell-lines with respect to miRCancer annotations. Ratio of expression of SF-miRNAs in MDA-MB-231 to MCF-10A (log scale). A total of 22 differentially expressed SF-miRNAs are listed with a minimal expression levels of >50 reads (sum of MCF-10A and MDA-MB-231 expression reads). The color signifies the comparison according to the consistency with miRCancer (green-consistent trend; grey stripes-not available; blue, mixed trend; red, opposite trend; white, minimal support (defined as miRNA with only 1-2 reported publications).

### 2.5. Inverse Expression Trend of SF-miRNAs and Total miRNAs from Breast Cancer Biopsies

The phenomenon of inverse directionality in expression of the SF-miRNAs in the breast cell-lines compared with the data collected from miRCancer is observed for 6 miRNAs (out of 22 listed SF-miRNAs). Still for another 9 of the most significant SF-miRNAs, miRCancer provides no or poorly supported information (<3 publications, Figure 6). This shortage in data prompted us to investigate the relative expression of these miRNAs from clinical samples. To this end, we applied Kaplan-Meier (KM)-plotter for the breast cancer miRNAs collection [63].

Figure 7 (top) shows expression plots of all top significant 22 SF-miRNAs (as in Figure 6) in the non-malignant MCF-10A cells and the metastatic MDA-MB-231 cells. The SF-miRNAs above the diagonal (Figure 7, top) are specified by higher expression in the metastatic cells (MDA-MB-231) versus the non-malignant cell (MCF-10A). The color of each SF-miRNA indicates the impact of the miRNA expression on patient survival as analyzed from two large breast cancer cohorts: the TCGA (1062 samples) [64] and METABRIC (1262 samples) [65]. The expression of each miRNA in each of the cohorts was split to high and low expressing quantiles. Figure 7 (bottom) shows representative miRNAs by their Kaplan-Meier survival plots. For each miRNA the hazard ratio (HR, 95% confidence intervals) and the logrank P-value were calculated. Information was available and collected for almost all SF-miRNAs (21 of 22; see Supplemental Table S6). The expression of SF-miR-21 is elevated in MDA-mB-231 cells compared to non-malignant MCF-10A, and is also in agreement to its malignancy index as tumorigenic (red), while the expression level of SF-miR-148a is decreased with malignancy also in agreement with its malignancy index as tumor suppressor (green). However, we found that for 5 SF-miRNAs whose expression level increased with the degree of malignancy (miR-100; let-7f-1; let-7f-2; let-7g; miR-30a) the clinical calculation of the KM survival plot shows that these miRNAs are associated with a hazard ratio (HR) below 1.0, which is consistent with a protective biomarker (colored green). Furthermore, for 8 SF-miRNAs whose expression level decreased with increased cellular malignancy (miR-6087; miR-1246; miR-7704; miR-3604; miR-622; miR-612; miR-5047; miR-7161), the clinical results are consistent with poor survival rate (colored red). Note that the HR and the logrank statistics for miR-622 and miR-7704 are exceptionally high (HR is 2.23). For the others SF-miRNAs (5), no significant difference is associated with the clinical survival rate (e.g. miR-221). Importantly, the majority of the strongly differentially expressed SF-miRNAs (13) show inconsistency between the expression trend in SF-miRNA and the clinical outcomes, suggesting that these miRNAs in the nucleus act on different targets. Only miR-21 and miR-222 are consistent with the role of miRNA as oncomiRs, and miR-148a as a tumor suppressor.

**Figure 7.**
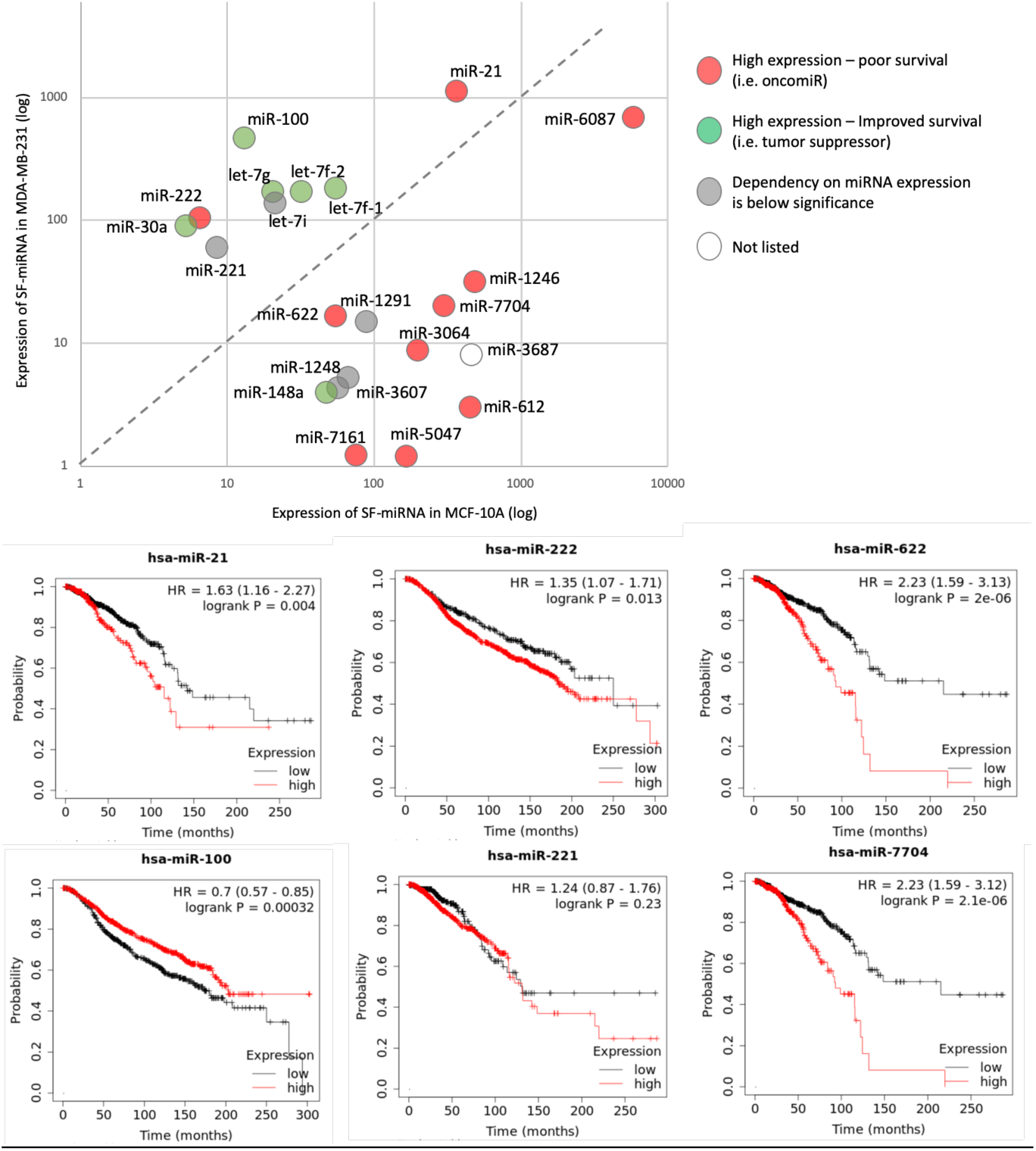
Expression of SF-miRNAs in different cell-lines vis-a-vis patient’s survival rate. (Top) Scatter plot of the SF-miRNAs expression in non-cancerous MCF-10A cells (x-axis) and metastatic MDA-MB-231 cells (y-axis) is presented. A total of 22 differentially expressed SF-miRNAs are listed with a minimal expression levels of >50 reads (sum of MCF-10A and MDA-MB-231 expression reads). The results of the Kaplan-Meier (KM) survival plots are color coded with miRNAs that are oncogenic (red), protective (green) and statistical insignificant (grey). Each tested miRNA is colored by the survival rate from breast cancer patients extracted from the TCGA (1062 samples) and METABRIC (1262 samples) projects. The KM survival plots analyze the survival (in months) associated with high and low expression quartiles of the selected miRNAs. Analysis is completed for 21 of the 22 SF-miRNAs (see Supplemental Table S6). (Bottom) Representative SF-miRNAs analyzed by KM survival plots with respect to the calculated hazard ratio (HR, 95% confidence intervals) and logrank with p-value <0.05.

### 2.6. Negative Correlation Between the Expression of SF-miR-7704 and the Oncogenic lncRNA HAGLR

The results presented in Figures 6 and 7 argue that the collection of SF-miRNAs is different from that of the cytosolic miRNAs (see also [54]). Furthermore, many of the spliceosome associated miRNAs were derived from intergenic regions (e.g. Table 1). Thus, the presence of miRNA sequences in the spliceosome suggests they carry overlooked function, likely different from the classical translation suppression that characterized cytosolic miRNAs. SF-miR-7704 was previously shown to negatively regulate the lncRNA HAGLR [54]. Here we analyzed the changes in expression of miR-7704 and HAGLR in the breast cancer cell-lines. The genomic position of miR-7704 with respect to the neighboring genes is illustrated in Figure 8A. We show that the level of expression of miR-7704 varies in each of the tested cell-lines with maximal expression in the MCF-10A and negligible expression in the MDA-MB-231 cells (Figure 8B). RT-PCR analyses of the expression of HAGLR and HOXD1 (Figures 8C, 8D) confirm the relative changes in expression across the breast cell-lines. Figures 8C and 8D demonstrate that the expression level of HAGLR is hardly noticeable in MCF-10A cells, but maximal expression is seen in the MDA-MB-231 cells. This high expression level of HAGLR is negatively correlated with the low expression level of miR-7704. These findings were repeated when RNA was extracted from the cells or from isolated nuclei. The expression level of HOXD1 in the three cell-lines remains low, and will not be further discussed.

**Figure 8.**
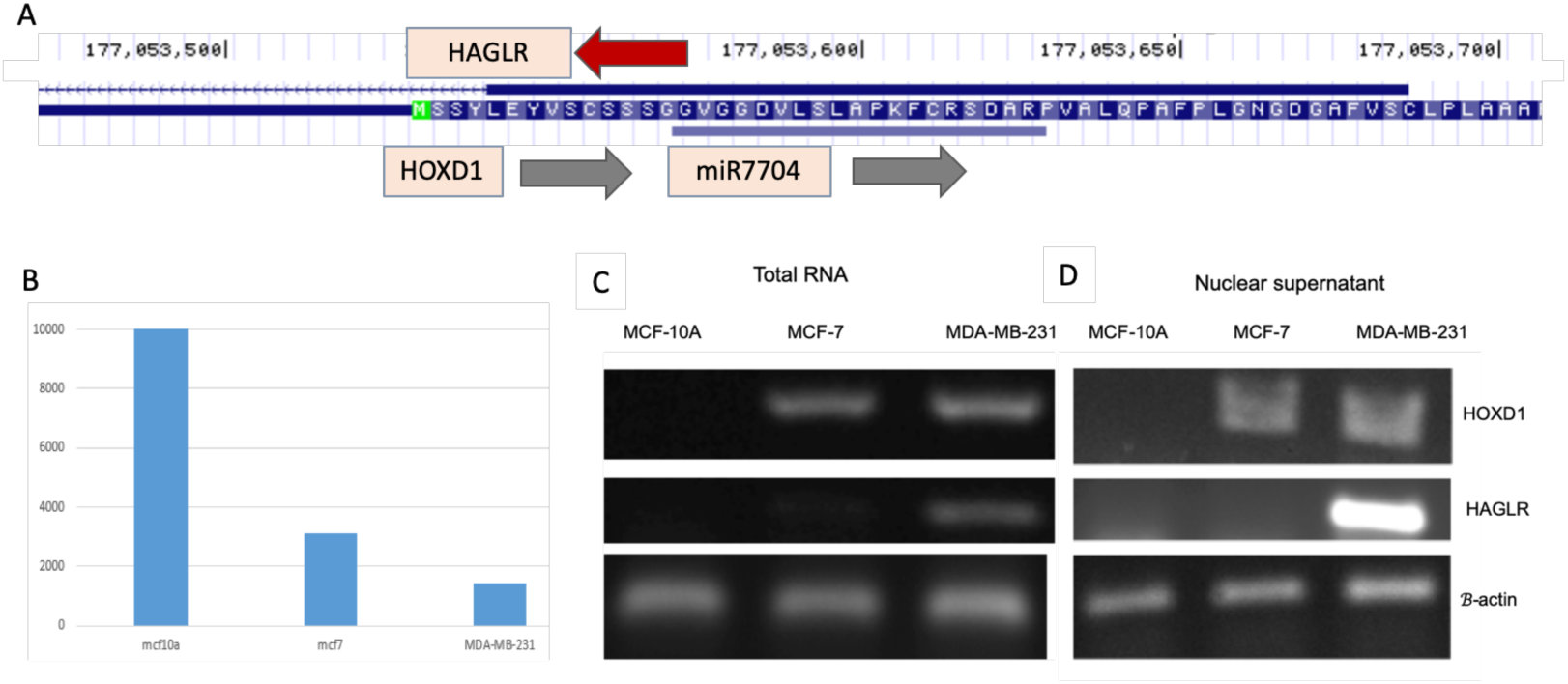
Negative correlation between the expression of miR-7704 and HAGLR. **(A)** Genome browser UCSC view for miR-7704 and the overlap with HOXD1 and the lncRNA HAGLR. **(B)** Average expression levels of miR-7704 from the SF based triplicates of MCF-10A, MCF-7 and MDA-MB-231 cells. (C, D) Comparison of the results of RT-PCR assays of HOXD1 and HAGLR expression as measured from total RNA **(C)** and nuclear RNA **(D)** in the three tested breast cell-lines. Results represent average of RT-PCR from 3 biological repeats. The identity of the extracted bands was confirmed by DNA sequencing

### 2.7. Manipulating the Expression Level of miR-7704 Dictates HAGLR Expression Level

To test the effect of miR-7704 on the expression of HAGLR we transfected (in triplicates) each of the three-breast cell-lines with anti-miR-7704 inhibitor. We next extracted nuclear RNA from each of the transfected cell-lines, and measured the expression levels of miR-7704 and HAGLR by quantitative-PCR. Cells transfected with non-relevant anti-miRNA and untreated cells were used as controls (see Materials and Methods). As shown in Figure 9A, transfection of MCF-10A with anti-miR-7704 inhibitor reduced the level of nuclear miR-7704 to 30% as compared with the non-treated cells, while a non-relevant antimiR had a negligible effect. The decrease in the level of nuclear miR-7704, is followed by >2 folds upregulation in the level of lncRNA HAGLR (Figure 9B, lower panel). When the same protocol was applied for MCF-7 cells (Figure 9B), transfection with anti-miR-7704 reduced the level of nuclear miR-7704 in the cells to 55% of its original level of non-treated cells. In these cells, the upregulation in level of HAGLR was ∼2 folds (Figure 9B, lower panel). Transfection of MDA-MB-231 cells with anti-miR-7704 reduced the level of miR-7704 to ∼20% and its original level, resulting in a 2-fold increase in the level of HAGLR (Figure 9C). We conclude that miR-7704 negatively regulates the expression of the oncogenic lncRNA HAGLR.

**Figure 9.**
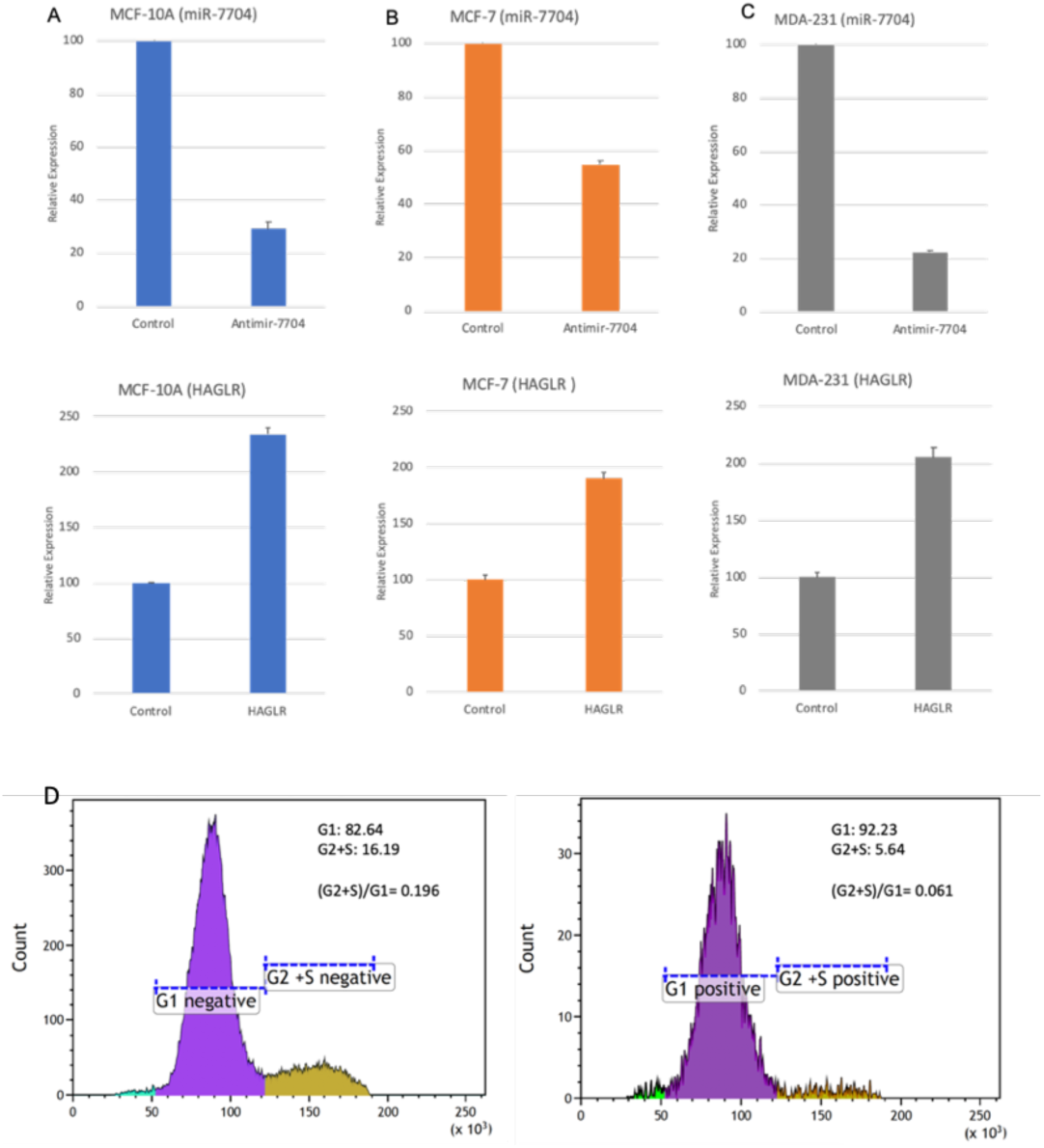
Inhibition of miR-7704 Elevated HAGLR Expression while Overexpression Reduced Cell Division Rate. (A-C) Inhibition of miR-7704 expression upregulates the expression of HAGLR. Real time PCR analysis of the effect of inhibition of miR-7704 on the nuclear expression of HAGLR. The results shown are from MCF-10A cells **(A)**, MCF-7 cells **(B)** and MDA-MB-231 cells **(C)**. Transfection with Anti-miR-7704 inhibitor resulted in down-regulation of miR-7704 (upper panel), and increase in the expression level of HAGLR mRNA (lower panel). The results of quantitative PCR of 3 independent biological preparations are shown. The expression levels of HAGLR was normalized to internal control of ß-actin expression from the same preparation. **(D)** The FACS analysis of the cells cycle following transfecting the MDA-MB-231 cells with miR-7704. The sorting is based on GFP expression (488nm) that indicates the parallel expression of miR-7704. Single live cells were stained with Hoechst to quantify the amounts of DNA in each cell. The data was partitioned to sub-Go/G1 (green), Go/G1 (marked G1, purple) and S/G2/M (marked G2+S, brown). The untransfected (left) and GFP positives (right) are representative results of 3 independent biological experiments (<15% variations). The fraction of G2+S fraction was reduced and occupied 5.6% instead of the 16.9% in the untransfected control (3.2 folds reduction). The reduced fraction of S/G2 relative to G0/G1 is used to assess the suppression in the cell division.

The manipulation of miR-7704 levels resulted in rebalance in the expression of HAGLR (Figures 9A-9C). We asked whether overexpression of miR-7704 in the metastatic cell MDA-MB-231 can lead to any attenuation in the metastatic characteristics of the cells. For this aim we analyzed the unsynchronized cells by quantifying the major stages of the cell cycle - the G0/G1 and the G2/S/M. A shift in the fraction occupied by the G2/S/M stage is indicative of a change in cell division. We compared these major stages in the cell cycle for the basal level of miR-7704, and following its overexpression (in biological triplicates). MDA-MB-231 cells were transfected with a plasmid expressing has-miR-7704 (see Materials and Methods), using an empty vector as control. As miR-7704 overexpression is accompanied by a parallel expression of GFP, we used the GFP fluorescence signal to sort cells that were transfected and contains elevated levels of miR-7704. Figure 9D shows the results from MDA-MB-231 cells and their cell cycle partition. The analysis shows a reduction in cell division as a result of miR-7704 overexpression. In all experiments there were 3000-5000 transfected cells and >30,000-50,000 cells in the untransfected control. The fraction of S/G2/M changed from16.2% to 6.1% of the entire tested cells. We conclude that the increase in the level of miR-7704 can be used to attenuate and alter the rate of cell division, and thus potentially to slow down the uncontrolled cell division that is a hallmark of cancer.

## 3. Discussion

miRNAs are mainly known for their role in inhibition of translation in the cytoplasm [3-7]. The finding of miRNAs in the nucleus [17-21] led to the discovery of novel functions for miRNA in the nucleus [22, 23]. Nuclear miRNAs are involved in the regulation of non-coding RNAs (ncRNAs), in transcriptional silencing, activation, and inhibition (Reviewed in 20, 21). However, the functions of miRNAs in the nucleus are not yet well understood, and require further studies to elucidate their full potential. One plausible hypothesis for the origin of the miRNA in the nucleus concerns the localization of many miRNA genes in introns. In this case, the biogenesis is likely linked to the endogenous spliceosome and accompanies the expression of the miRNA host genes. However, we have shown that many of the spliceosomal miRNAs are not embedded in introns [54, 55]. While previous studies of nuclear miRNAs analyzed the whole nuclear miRNA population, our study focuses on miRNAs enriched in the spliceosomal fraction (SF). The identification of a large collection of SF-miRNAs derived sequences in HeLa cells [54], and the reported deregulation of miRNAs in cancer [1], and particularly in breast cancer [2, 66] have directed us to characterize SF-miRNA in established breast cancer cell-lines.

Female breast cancer is the second most commonly diagnosed cancer and the fifth leading cause of cancer-related death worldwide [2, 67-70]. Oncogenic and tumor suppressor miRNAs were shown to affect cell proliferation, apoptotic response, and metastasis in breast cancer [2]. However, the potential of SF-miRNAs to act as breast cancer indicators and as a source for novel therapeutic targets has not been studied before and is of utmost importance.

Sequencing of the SF small RNA from three breast-derived cell-lines revealed a rich collection of miRNA derived sequences that signified each of the tested cells and indicated the profile along the cancer progression from MCF-10A, to the metastatic MDA-MB-231 cells. Among the SF-miRNA we identified numerous known cancer-associated oncogenic miRNAs (e.g. miR-21, miR-221, miR-222) and tumor suppressor miRNAs (e.g. miR-100, miR-612, miR-30a and several members of the let-7 family; Table 1). Importantly, we found significant changes in the expression of SF-miRNAs with increased malignancy of these cells. These changes are portrayed in variations in SF-miRNA types, their level of expression (Table 1, Figures 2-3, 5), but also in the composition of the segmental regions of expressed SF-miRNAs (Figure 4). The SF-miRNA sequences are not only confined to mature miRNAs. Actually, only ∼25% of SF-miRNAs are made up of mature miRNAs in non-malignant MCF-10A cells, but this fraction increases to 60% and 68% in MCF-7 and MDA-MBB-231, respectively (Figure 4B). Other changes are probably underlying the processing of the miRNA related sequences as reflected by difference in length distribution in each cell-line (Figures 4C).

Notably, several SF-miRNAs show large changes in their level of expression across the 3 cell-lines (Figure 5). Several miRNAs increase in expression along with the cellular malignant state, but another set of SF-miRNAs exert the opposite trend with decreased expression with cells specified by high malignancy. Importantly, the changes that we observe in expression level of SF-miRNAs in specific cases contradict with miRCancer literature reports. The changes in expression level of SF-miRNAs between the non-malignant breast cells MCF-10A and the metastatic breast cell-line MDA-MB-231 are compared to those reported in cancerous sample versus healthy samples (from miRCancer). Recall that the reports in miRCancer rely on a cellular cytosolic view (Figure 6). We found that tumorigenic miR-21, miR-221 and miR-222, as well as tumor suppressor miR-148a and miR-622 show the same trend in both cases. However, for the known tumor suppressor set of let-7g, let-7i, let-7f-2, let-7f-1, miR-30a and miR-100, the SF-miRNAs show an opposite trend and were signified by an increase in expression in breast cancer. Also, miR-1246 that is known as tumorigenic associated miRNA, its expression in the spliceosome is suppressed in cells that resemble cancer of high malignancy. An extended analysis performed on breast cancer biopsies of very large cohorts (>1000 samples each) [64, 65] confirmed and substantiated the discrepancy between the expression trend of some SF-miRNAs and the observation reported from the literature (based on miRCancer). We noted that the majority of the SF-miRNAs shows inverse expression trend in view of total miRNA analysis (13 of 21 informative miRNAs). The others are either below statistical significance and only 3 miRNAs are consistent with the clinical findings (Figure 7). We thus postulate that the majority of significant and highly expressed SF-miRNAs act on alternative targets in the spliceosome, leading to an inverse impact on malignancy. Specifically, several known oncogenic miRNAs act as tumor suppressors in the spliceosome while several tumor suppressors miRNAs are tumorigenic in the spliceosome. Thus, further study might reveal overlooked nuclear targets that can be attractive lead for drug design of breast cancer. Despite the partial overlap in the information provided from miRCancer and the breast cancer survival results we observed high degree of agreement, where miR-100, miR-30a and several of the let-7 members show the same trend, which is opposite to that of the SF-miRNAs.

The presence of SF-miRNAs suggests their potential nuclear functions. Bioinformatic analysis for targets of SF-miRNAs through base-pairing, highlighted the potential of such sequences to regulate gene expression and splicing [54]. Here we focused on miR-7704, which is complementary to sequences of the first exon of the lncRNA HAGLR (Figure 8). We showed that SF-miR-7704 decreases in expression level with the severity of breast cancer (Table 1, Figure 8) and it is negatively correlated with the expression of HAGLR. We further demonstrated that manipulating cells by suppressing the expression level of miR-7704 negatively regulates HAGLR expression (Figure 8). These observations corroborate the findings made in HeLa cells [54]. It should be noted that according to the clinical calculation of the KM survival plot miR-7704 is associated with exceptionally high HR and logrank statistics which is consistent with its role as a tumorigenic function (Figure 7, Supplemental Table S6). Yet, the level of miR-7704 is decreased with tumorigenicity and it governs the expression level of HAGLR. Notably, HAGLR is a lncRNA that was implicated in many cancers and was shown to be upregulated in different human cancers including bladder, cervical, colorectal, gastric, ovarian, prostate cancers, glioma, hepatocellular carcinoma, melanoma, osteosarcoma and non-small cell lung cancer [56]. HAGLR has been shown to play a role in the development and progression of these cancers and its expression level was correlated with the clinical features of cancer. Although the presence of HAGLR was not previously reported in breast cancer [56], we demonstrate here that the level of HAGLR is positively correlated with the severity of cancer, with the highest expression level found in the metastatic breast cancer cells MDA-MB-231. We have thus identified HAGLR as the nuclear target of SF-miR-7704, which acts as a tumor suppressor in the spliceosomal context. The negative regulation of HAGLR by miR-7704 can serves as an overlooked path in controlling aggressive breast cancer.

## 4. Materials and Methods

### 4.1. Plasmids

For overexpression of hsa-mir-7704, the pre-miR-7704 sequence (5’-CGGGGTCGGCGGCGACGTGCTCAGCTTGGCACCCAAGTTCTGCCGCTC CGACGCCCGGC-3’) was cloned into the pPRIME-CMV-GFP-FF3 vector at the Xho1 and EcoR1 sites by GENEWIZ (South Plainfield, NJ, USA), generating pPRIME-hsa-mir-7704-CMV-GFP-FF3 plasmid [54]. A pPRIME-CMV-GFP-FF3 empty vector was used as a control.

### 4.2. Cells

Spliceosomes and RNA were isolated from the following cell-lines: MCF-10A, non-tumorigenic human mammary epithelial cell-line (ATCC, CRL-10317); MCF-7 mammary epithelial adenocarcinoma cell-line (ATCC HTB-22); MDA-MB-231 aggressive metastatic tumorigenic human mammary epithelial cell-line (ATCC HTB-26).

### 4.3. Isolation of Supraspliceosomes

All isolation steps were conducted at 4°C. Supraspliceosomes were prepared from nuclear supernatants enriched in supraspliceosomes as previously described [51] from the following cell-lines: MCF-10A, MCF-7 and MDA-MB-231. Briefly, nuclear supernatants were prepared from purified cell nuclei by microsonication of the nuclei and precipitation of the chromatin in the presence of excess of tRNAs. The nuclear supernatant was fractionated on 10-45% (vol/vol) glycerol gradients. Centrifugations were carried out at 4°C in an SW41 rotor run at 41 krpm for 90 min [or an equivalent @2t = 2500 (@ is in krpm; t is in hr)]. The gradients were calibrated using the tobacco mosaic virus as a 200S sedimentation marker. Supraspliceosome peak fractions were confirmed by Western Blot and by electron microscopy visualization.

### 4.4. Protein Detection

Western blotting (WB) analyses were performed as previously described [59]. We used anti-hnRNP G (kindly provided by Prof. Stefan Stamm, University of Kentucky, Lexington), visualized with horseradish peroxidase conjugated to affinity-pure Goat anti-Rabbit IgG (H+L; Jackson Immunoreaserch, 1:5000). The MAb-104 that is directed against the phosphorylated epitopes of the SR proteins [71] was used as previously described [72].

### 4.5. RNA Isolation from Supraspliceosomes and Deep Sequencing

RNA was extracted as previously described [51] from supraspliceosomes prepared from each of the different three breast cell-lines: the two breast cancer cells at different stages of malignancy: MCF-7 and MDA-MB-231; and the non-tumorigenic breast cell-line MCF-10A. The integrity of the RNA was evaluated by an Agilent 2100 bioAnalyzer. For small RNA library construction, ∼1 µg of RNA was used. After phosphatase and T4 polynucleotide kinase (PNK) treatments, the RNA was ethanol precipitated to enrich for small RNA, and small RNA libraries (in triplicates) were prepared according to NEBNext Small RNA Library Prep Set for Illumina (Multiplex Compatible) Library Preparation Manual. Adaptors were then ligated to the 5’ and 3’ ends of the RNA, and cDNA was prepared from the ligated RNA and amplified to prepare the sequencing library. The amplified sequences were purified on E-Gel® EX 4% Agarose gels (ThermoFisher # G401004), and sequences representing RNA smaller than 200 nt were extracted from the gel. The library was sequenced using the Illumina NextSeq 500 Analyzer. The sequencing data, after removal the adaptors and filtering out low quality sequences, were aligned to mirBase (Release 21). In addition, the filtered high-quality fragments were mapped to the human transcriptome of hg19 gtf file from UCSC provided by Galaxy. The hg19 transcriptome contains 963,559 exons from 45,314 transcripts. Sequences aligned to the miRNA genes as compiled in miRBAse are reported.

### 4.6. Next Generation Sequencing (NGS) Analysis

RNA was extracted from three independent biological preparations from the supraspliceosome fractions from each of the different breast cell-lines. NGS was performed for each sample on small RNA (<200 nt) molecules using standard Illumina Protocol. Each library consisted of an average of 18.5M (±9 M) reads of maximum length 76 (see supplementary **Table S1**).

Raw data of the sequenced small RNA were trimmed using Cutadapt ver. 1.13. Low-quality reads were filtered out using FASTX toolkit. Reads from the SF were aligned against human genome hg19 and miRbase database (version 21) using TopHat 2.1.1, allowing 90% sequence identity and a maximum of two mismatches. Reads whose start or end position were mapped to miRNA genes were considered. High quality reads from the three SF preparations of each cell-line were combined. Out of the mapped reads, only reads of length ≥17 were considered. miRNA gene aligned sequences refer to all mapped, high quality reads that are aligned to any of the pre-miRNA as defined by miRBase. For the rest of the analyses, only miRNA aligned sequence with ≥10 reads per cell-line were considered. This threshold was used to ensure reliable support in view of the limited total reads assigned from supraspliceosomes.

### 4.7. Validation of Gene Expression

#### 4.7.1. RT-PCR

RT-PCR was performed on RNA extracted from the cell-lines described above, and from nuclear supernatants of the above cells as described [60]. The following sets of primers for HAGLR was used: Forward (exon 1) 5’-CGTCGGAGCGGCAGAACTT-3’ and Reverse 5’-AAGGGCCCATTTTCAGGCCA-3’ (exon 2). The primers for HOXD1 are: Forward (exon 1) 5’-ATTTACCTCCGGCTCACTCG-3’ and Reverse: 5’-AGGTGCAAGCAGTTGGCTAT-3’ (exon 2). The identity of all PCR products was confirmed by sequencing. Each experiment was repeated at least 3 times. The relative abundance was quantified in view of the intensity of the ß-actin that was used as a control. The ß-actin Forward and Reverse primers, for an amplicon of 140 nt, are: 5’-CTGGAACGGTGAAGGTGACA-3’ and 5’-AAGGGACTTCCTGTAACAATGCA-3’, respectively.

#### 4.7.2. Transfection and RNA Isolation

MCF-10A, MCF-7 and MDA-MB-231 cells were each grown in six-well plates. For downregulation of hsa-mir-7704, the cells were transfected with Anti-hsa-mir-7704 inhibitor AM29132 (ThermoFisher) according to manufacturer’s instructions at 100 nM for 48 hr. As controls we used the same cells transfected with mirVana miRNA inhibitor, negative control #1 (Ambion), and non-treated cells.

Nuclear RNA isolation was performed as previously described [44]. Briefly, 48 hr post transfection, the six-well plates were washed with PBS followed by the addition of 175 µl of cold RLN buffer (50 mM Tris pH 8, 140 mM NaCl, 1.5 mM MgCl_2_ and 0.5% NP40). The cells were then scraped and moved to an Eppendorf tube on ice for 5 min. Centrifugation for 2 min at 300 g, at 4oC, was then performed. The supernatant was transferred to a new tube and the pellet (nuclei) was centrifuged again. RNA was then extracted from nuclei with miRNeasy mini kit (Qiagen), following the manufacturer’s instructions. All experiments were performed with at least three biological replicates. For overexpression of miR-7704 the MDA-MB-231 cells were grown in six-well plates and transfected for 48h with pPRIME-miR-7704-CMV-GFP-FF3 plasmid (5.5 µg/well) using Lipofectamine-2000 reagent (ThermoFisher). As control, we used transfection with 5.5 µg/well of the empty vector. Removal of adherent cells from the plates was according to routine protocol (using Trypsin, and PBS with 10% FCS to inactivate it). Cell suspension was transferred to the FACS tube and stained with 5 µl Hoechst (1 mg/ml, Sigma #33342) as a measure for DNA cell content. Cell cycle analysis was performed using a fluorescence analysis (using FACS BD-Facsaria III Bactlab) equipped with 405 nm and 488 nm lasers, for Hoechst and GFP, respectively. Gating applied for all live cells in the preparation. FACS data analysis used Kaluza and the Beckman Coulter Flow Cytometry Analysis Software.

### 4.8. Quantitative PCR

#### 4.8.1. TaqMan microRNA Assay

For RT-PCR of miR-7704, the TaqMan Advanced miRNA cDNA synthesis kit (ThermoFisher) was used according to the manufacturer’s instructions, which included polyA tailing, adaptor ligation and miR-Amp reaction as previously described [54].

#### 4.8.2. RT of mRNA

RT of nuclear RNA was performed using the High Capacity cDNA Reverse Transcription Kit (ThermoFisher) according to the manufacturer’s instructions, using RT Random Primers, and MultiScribeTM Reverse Transcriptase.

#### 4.8.3. Quantitative PCR reaction

mRNA and miR-7704 levels were measured using the TaqMan Fast Advanced Mix (ThermoFisher) and the following TaqMan Assays with FAM/MGB-NFQ primers/probe: TaqMan Advance MiR Assay hsa-mir-7704, 480576_mir (AB-25576 ThermoFisher); TaqMan Gene Expression Assays MTO, XS/PC: beta actin: Hs99999903_m1 (AB-4453320); TaqMan Gene Expression Assays /PC: HOXD1: Hs04334671_g1 (AB-4448892); Custom TaqMan Copy Number Assays, SM/PC: HAGLR_NR_033979.2 (AB-4400294). Fw primer: TGCCAAGCTGAGCACGTC, Rev: TACTCCAGATCTGGGGAC, FAM Probe: ACGTACTCCAGATCTG. Assays were performed according the manufacturer’s instructions. Amplification was carried out using a QuantStudio 12K Flex Real-Time PCR System (for downregulation), and StepOnePlus Real-Time PCR System (for overexpression), for 40 cycles at annealing temperature of 60°C. Analysis was performed using the delta-delta C_T_, 2-ΔΔCT method. All experiments were performed in at least three biological repeats.

## 5. Conclusions

Comparison of the miRNA sequences associated with the endogenous spliceosome of human breast-derived cell-lines that range in their cancerous states (from the non-malignant MCF-10A, the malignant MCF-7, and the highly metastatic MDA-MB-231 cells), revealed changes with malignancy of the rich collection of SF-miRNAs, including changes in the miRNA types, level of expression, and the composition of pre-miRNA segmental regions.

Notably, a large fraction of the abundant SF-miRNAs (e.g. miR-100, miR-30a, and let-7 family members) show an opposite consequence on the expression trend when compared to the literature and breast cancer clinical samples. These findings indicate that SF-miRNA expression consequence on cancer is opposite to that of the cellular miRNAs. It further implies that these SF-miRNAs act on different, yet unexplored, targets in the nucleus compared to their cytoplasmic ones. One such miRNA is miR-7704 whose genomic position overlaps HAGLR, a cancer-related lncRNA. In the three tested breast cell-lines, we quantified an inverse expression of miR-7704 with respect to HAGLR. Moreover, inhibiting miR-7704 expression led to an increased level of HAGLR, while its overexpression led to a reduction in cell-division rate in MDA-MB-231 cells. While miR-7704 acts as oncomiR in breast cancer patients, it has a tumor suppressing function in the SF, with HAGLR being its nuclear target. The negative regulation of HAGLR by miR-7704 can serves as an overlooked path in controlling aggressive breast cancer. Altogether, we highlight miRNAs in the spliceosome as an unexplored route for breast cancer therapeutics.

## Supplementary Materials

**Figure S1**. Spearman rank correlation between all 9 samples of the SF-miRNA collection identified in each sample.

**Table S1**. A statistical summary of the sequencing of breast cell-lines libraries.

**Table S2**. Amounts of aligned reads from SF-miRNAs (raw data) associated with each of the nine sample preparations from three cell-lines. Source data for **Figure** 2 and **Supplemental Figure S1.**

**Table S3**. A list of SF-miRNAs according to their statistical significance following normalization by DESeq2. Source for **Figure 3, Table** 1 and **Figure 5**.

**Table S4**. List of SF-miRNA according to segmental regions of a prototype of pre-miRNAs for each of the tested cell-lines. A summary for the segmental region partition for cell-lines. Source data for **Figure 4**.

**Table S5**. Comparison of changes of 22 SF-miRNAs in the three-breast cell-lines with data from literature (miRCancer). Source data for **Figure 6**.

**Table S6**. Comparison of changes of 22 SF-miRNAs in the three-breast cell-lines with breast cancer KM survival plots. Source data for **Figure 7**.

## Author Contributions

R.S. and M.L. conceived and designed the study. A.R.P., K.Z., Y.C., and T.E. performed the experiments; R.S., and M.L. designed the RNA-Seq experiments. M.L. and S.M.A. designed the bioinformatic analyses. S.M.A. performed the data analysis, bioinformatic and statistical analyses, of the RNA-Seq data; R.S. and M.L. wrote the paper. All authors discussed the findings, and contributed to the final manuscript.

## Acknowledgments

We would like to thank Aviva Petcho for excellent technical assistance. We thank the system team of the Computer Science and Engineering at the Hebrew University for their support, and Prof. Stefan Stamm (University of Kentucky) for the anti-hnRNP G antibodies.

## References

1. Di Leva G, Garofalo M, Croce CM: microRNA in cancer. Annu Rev Pathol 2014, 9:287-314.

2. Loh HY, Norman BP, Lai KS, Rahman N, Alitheen NBM, Osman MA: The Regulatory Role of MicroRNAs in Breast Cancer. Int J Mol Sci 2019, 20:4940.

3. Bartel DP: MicroRNAs: target recognition and regulatory functions. Cell 2009, 136:215–233.

4. Fabian MR, Sonenberg N: The mechanics of miRNA-mediated gene silencing: a look under the hood of miRISC. Nat Struct Mol Biol 2012, 19:586–593.

5. Krol J, Loedige I, Filipowicz W: The widespread regulation of microRNA biogenesis, function and decay. Nature reviews Genetics 2010, 11:597–610.

6. Ha M, Kim VN: Regulation of microRNA biogenesis. Nat Rev Mol Cell Biol 2014, 15:509–524.

7. Bartel DP: Metazoan MicroRNAs. Cell 2018, 173:20–51.

8. Gregory RI, Yan KP, Amuthan G, Chendrimada T, Doratotaj B, Cooch N, Shiekhattar R: The Microprocessor complex mediates the genesis of microRNAs. Nature 2004, 432:235–240.

9. Han J, Lee Y, Yeom KH, Kim YK, Jin H, Kim VN: The Drosha-DGCR8 complex in primary microRNA processing. Genes Dev 2004, 18:3016–3027.

10. Han J, Lee Y, Yeom KH, Nam JW, Heo I, Rhee JK, Sohn SY, Cho Y, Zhang BT, Kim VN: Molecular basis for the recognition of primary microRNAs by the Drosha-DGCR8 complex. Cell 2006, 125:887–901.

11. Landthaler M, Yalcin A, Tuschl T: The human DiGeorge syndrome critical region gene 8 and Its D. melanogaster homolog are required for miRNA biogenesis. Curr Biol 2004, 14:2162–2167.

12. Lee Y, Ahn C, Han J, Choi H, Kim J, Yim J, Lee J, Provost P, Radmark O, Kim S, Kim VN: The nuclear RNase III Drosha initiates microRNA processing. Nature 2003, 425:415–419.

13. Yi R, Qin Y, Macara IG, Cullen BR: Exportin-5 mediates the nuclear export of pre-microRNAs and short hairpin RNAs. Genes Dev 2003, 17:3011–3016.

14. Kim YK, Kim VN: Processing of intronic microRNAs. Embo J 2007, 26:775–783.

15. Kozomara A, Birgaoanu M, Griffiths-Jones S: miRBase: from microRNA sequences to function. Nucleic Acids Res 2019, 47:D155–D162.

16. Acunzo M, Romano G, Wernicke D, Croce CM: microRNA and cancer - a brief overview. Advances in Biological Regulation 2015, 57:1–9.

17. Roberts TC: The MicroRNA Biology of the Mammalian Nucleus. Mol Ther Nucleic Acids 2014, 3:e188.

18. Guil S, Esteller M: RNA-RNA interactions in gene regulation: the coding and noncoding players. Trends Biochem Sci 2015, 40:248–256.

19. Huang V, Li LC: miRNA goes nuclear. RNA Biol 2012, 9:269–273.

20. Liu H, Lei C, He Q, Pan Z, Xiao D, Tao Y: Nuclear functions of mammalian MicroRNAs in gene regulation, immunity and cancer. Mol Cancer 2018, 17:64.

21. Catalanotto C, Cogoni C, Zardo G: MicroRNA in Control of Gene Expression: An Overview of Nuclear Functions. Int J Mol Sci 2016, 17.

22. Liao JY, Ma LM, Guo YH, Zhang YC, Zhou H, Shao P, Chen YQ, Qu LH: Deep sequencing of human nuclear and cytoplasmic small RNAs reveals an unexpectedly complex subcellular distribution of miRNAs and tRNA 3’ trailers. PloS one 2010, 5:e10563.

23. Jeffries CD, Fried HM, Perkins DO: Nuclear and cytoplasmic localization of neural stem cell microRNAs. Rna 2011, 17:675–686.

24. Ohrt T, Mutze J, Staroske W, Weinmann L, Hock J, Crell K, Meister G, Schwille P: Fluorescence correlation spectroscopy and fluorescence cross-correlation spectroscopy reveal the cytoplasmic origination of loaded nuclear RISC in vivo in human cells. Nucleic Acids Res 2008, 36:6439–6449.

25. Ameyar-Zazoua M, Rachez C, Souidi M, Robin P, Fritsch L, Young R, Morozova N, Fenouil R, Descostes N, Andrau JC, et al: Argonaute proteins couple chromatin silencing to alternative splicing. Nat Struct Mol Biol 2012, 19:998–1004.

26. Fasanaro P, Greco S, Lorenzi M, Pescatori M, Brioschi M, Kulshreshtha R, Banfi C, Stubbs A, Calin GA, Ivan M, et al: An integrated approach for experimental target identification of hypoxia-induced miR-210. J Biol Chem 2009, 284:35134–35143.

27. Leucci E, Patella F, Waage J, Holmstrom K, Lindow M, Porse B, Kauppinen S, Lund AH: microRNA-9 targets the long non-coding RNA MALAT1 for degradation in the nucleus. Sci Rep 2013, 3:2535.

28. Hansen TB, Wiklund ED, Bramsen JB, Villadsen SB, Statham AL, Clark SJ, Kjems J: miRNA-dependent gene silencing involving Ago2-mediated cleavage of a circular antisense RNA. EMBO J 2011, 30:4414–4422.

29. Tang R, Li L, Zhu D, Hou D, Cao T, Gu H, Zhang J, Chen J, Zhang CY, Zen K: Mouse miRNA-709 directly regulates miRNA-15a/16-1 biogenesis at the posttranscriptional level in the nucleus: evidence for a microRNA hierarchy system. Cell Res 2012, 22:504–515.

30. Nishi K, Nishi A, Nagasawa T, Ui-Tei K: Human TNRC6A is an Argonaute-navigator protein for microRNA-mediated gene silencing in the nucleus. RNA 2013, 19:17–35.

31. Kim DH, Saetrom P, Snove O, Jr., Rossi JJ: MicroRNA-directed transcriptional gene silencing in mammalian cells. Proceedings of the National Academy of Sciences of the United States of America 2008, 105:16230–16235.

32. Younger ST, Pertsemlidis A, Corey DR: Predicting potential miRNA target sites within gene promoters. Bioorg Med Chem Lett 2009, 19:3791–3794.

33. Morris KV, Santoso S, Turner AM, Pastori C, Hawkins PG: Bidirectional transcription directs both transcriptional gene activation and suppression in human cells. PLoS genetics 2008, 4:e1000258.

34. Han J, Kim D, Morris KV: Promoter-associated RNA is required for RNA-directed transcriptional gene silencing in human cells. Proc Natl Acad Sci U S A 2007, 104:12422–12427.

35. Schwartz JC, Younger ST, Nguyen NB, Hardy DB, Monia BP, Corey DR, Janowski BA: Antisense transcripts are targets for activating small RNAs. Nat Struct Mol Biol 2008, 15:842–848.

36. Yue X, Schwartz JC, Chu Y, Younger ST, Gagnon KT, Elbashir S, Janowski BA, Corey DR: Transcriptional regulation by small RNAs at sequences downstream from 3’ gene termini. Nat Chem Biol 2010, 6:621–629.

37. Chi SW, Zang JB, Mele A, Darnell RB: Argonaute HITS-CLIP decodes microRNA-mRNA interaction maps. Nature 2009, 460:479–486.

38. Shomron N, Levy C: MicroRNA-biogenesis and Pre-mRNA splicing crosstalk. J Biomed Biotechnol 2009, 2009:594678.

39. Mattioli C, Pianigiani G, Pagani F: Cross talk between spliceosome and microprocessor defines the fate of pre-mRNA. Wiley interdisciplinary reviews RNA 2014, 5:647–658.

40. Kataoka N, Fujita M, Ohno M: Functional association of the Microprocessor complex with the spliceosome. Mol Cell Biol 2009, 29:3243–3254.

41. Shiohama A, Sasaki T, Noda S, Minoshima S, Shimizu N: Nucleolar localization of DGCR8 and identification of eleven DGCR8-associated proteins. Exp Cell Res 2007, 313:4196–4207.

42. Janas MM, Khaled M, Schubert S, Bernstein JG, Golan D, Veguilla RA, Fisher DE, Shomron N, Levy C, Novina CD: Feed-forward microprocessing and splicing activities at a microRNA-containing intron. PLoS Genet 2011, 7:e1002330.

43. Allo M, Buggiano V, Fededa JP, Petrillo E, Schor I, de la Mata M, Agirre E, Plass M, Eyras E, Elela SA, et al: Control of alternative splicing through siRNA-mediated transcriptional gene silencing. Nat Struct Mol Biol 2009, 16:717–724.

44. Agranat-Tamir L, Shomron N, Sperling J, Sperling R: Interplay between pre-mRNA splicing and microRNA biogenesis within the supraspliceosome. Nucl Acids Res 2014, 42:4640–4651.

45. Papasaikas P, Valcarcel J: The Spliceosome: The Ultimate RNA Chaperone and Sculptor. Trends in biochemical sciences 2016, 41:33–45.

46. Kelemen O, Convertini P, Zhang Z, Wen Y, Shen M, Falaleeva M, Stamm S: Function of alternative splicing. Gene 2013, 514:1–30.

47. Lee Y, Rio DC: Mechanisms and Regulation of Alternative Pre-mRNA Splicing. Annual review of biochemistry 2015, 84:291–323.

48. El Marabti E, Younis I: The Cancer Spliceome: Reprograming of Alternative Splicing in Cancer. Front Mol Biosci 2018, 5:80.

49. Sperling J, Azubel M, Sperling R: Structure and Function of the Pre-mRNA Splicing Machine. Structure 2008, 16:1605–1615.

50. Sperling R: The nuts and bolts of the endogenous spliceosome. Wiley interdisciplinary reviews RNA 2017, 8:e1377.

51. Azubel M, Habib N, Sperling J, Sperling R: Native spliceosomes assemble with pre-mRNA to form supraspliceosomes. J Mol Biol 2006, 356:955–966.

52. Cohen-Krausz S, Sperling R, Sperling J: Exploring the architecture of the intact supraspliceosome using electron microscopy. J Mol Biol 2007, 368:319–327.

53. Zhang Z, Falaleeva M, Agranat-Tamur L, Pages AP, Eyras E E., Sperling J, Sperling R, Stamm S: The 5’ untranslated region of the serotonin receptor 2C pre-mRNA generates miRNAs and is expressed in non-neuronal cells. Exp Brain Res 2013, 230:387–394.

54. Mahlab-Aviv S, Boulos A, Peretz AR, Eliyahu T, Carmel L, Sperling R, Linial M: Small RNA sequences derived from pre-microRNAs in the supraspliceosome. Nucleic Acids Res 2018, 46:11014–11029.

55. Sperling R: Small non-coding RNA within the endogenous spliceosome and alternative splicing regulation. Biochim Biophys Acta Gene Regul Mech 2019, 1862 194406.

56. Li L, Wang Y, Zhang X, Huang Q, Diao Y, Yin H, Liu H: Long non-coding RNA HOXD-AS1 in cancer. Clinica chimica acta; international journal of clinical chemistry 2018, 487:197–201.

57. Spann P, Feinerman M, Sperling J, Sperling R: Isolation and visualization of large compact ribonucleoprotein particles of specific nuclear RNAs. Proc Natl Acad Sci USA 1989, 86:466–470.

58. Heinrich B, Zhang Z, Raitskin O, Hiller M, Benderska N, Hartmann AM, Bracco L, Elliott D, Ben-Ari S, Soreq H, et al: Heterogeneous Nuclear Ribonucleoprotein G Regulates Splice Site Selection by Binding to CC(A/C)-rich Regions in Pre-mRNA. J Biol Chem 2009, 284:14303–14315.

59. Raitskin O, Angenitzki M, Sperling J, Sperling R: Large nuclear RNP particles-the nuclear pre-mRNA processing machine. J Struct Biol 2002, 140:123–130.

60. Kotzer-Nevo H, de Lima Alves F, Rappsilber J, Sperling J, Sperling R: Supraspliceosomes at Defined Functional States Present portray the Pre-Assembled Nature of the pre-mRNA Processing Machine in the Cell Nucleus. Int J Mol Sci 2014, 15:11637–11664.

61. Falaleeva M, Pages A, Matsuzek Z, Hidmi S, Agranat-Tamir L, Korotkov K, Nevo Y, Eyras E, Sperling R, Stamm S: dual function of C/d box snoRNAs in rRNA modification and alternative pre-mRNA splicing. Proceedings of the National Academy of Sciences of the United States of America 2016, 113 Pages E1625-34.

62. Xie B, Ding Q, Han H, Wu D: miRCancer: a microRNA-cancer association database constructed by text mining on literature. Bioinformatics 2013, 29:638–644.

63. Nagy A, Lanczky A, Menyhart O, Gyorffy B: Validation of miRNA prognostic power in hepatocellular carcinoma using expression data of independent datasets. Sci Rep 2018, 8:9227.

64. Tomczak K, Czerwinska P, Wiznerowicz M: The Cancer Genome Atlas (TCGA): an immeasurable source of knowledge. Contemp Oncol (Pozn) 2015, 19:A68–77.

65. Mukherjee A, Russell R, Chin SF, Liu B, Rueda OM, Ali HR, Turashvili G, Mahler-Araujo B, Ellis IO, Aparicio S, et al: Associations between genomic stratification of breast cancer and centrally reviewed tumour pathology in the METABRIC cohort. NPJ Breast Cancer 2018, 4:5.

66. Emmanuel KN, Zacharias F, Valentinos P, Sofia K, Georgios D, Nikolaos KJ: The Impact of microRNAs in Breast Cancer Angiogenesis and Progression. Microrna 2019, 8:101–109.

67. Polyak K: Heterogeneity in breast cancer. J Clin Invest 2011, 121:3786–3788.

68. Rojas K, Stuckey A: Breast Cancer Epidemiology and Risk Factors. Clin Obstet Gynecol 2016, 59:651–672.

69. Fouad YA, Aanei C: Revisiting the hallmarks of cancer. Am J Cancer Res 2017, 7:1016–1036.

70. Feng Y, Spezia M, Huang S, Yuan C, Zeng Z, Zhang L, Ji X, Liu W, Huang B, Luo W, et al: Breast cancer development and progression: Risk factors, cancer stem cells, signaling pathways, genomics, and molecular pathogenesis. Genes Dis 2018, 5:77–106.

71. Roth MB, Murphy C, Gall JG: A monoclonal antibody that recognizes a phosphorylated epitope stains lampbrush chromosome loops and small granules in the amphibian germinal vesicle. JCB 1990, 111:2217–2223.

72. Yitzhaki S, Miriami E, Sperling J, Sperling R: Phosphorylated Ser/Arg-rich proteins: Limiting factors in the assembly of 200S large nuclear ribonucleoprotein particles. Proc Natl Acad Sci USA 1996, 93:8830–8835

