## Supplemental Materials for "Spliceosome-Associated MicroRNAs Identified in Breast Cancer Cells Act on Nuclear Targets and Are Potential Indicators for Tumorigenicity"

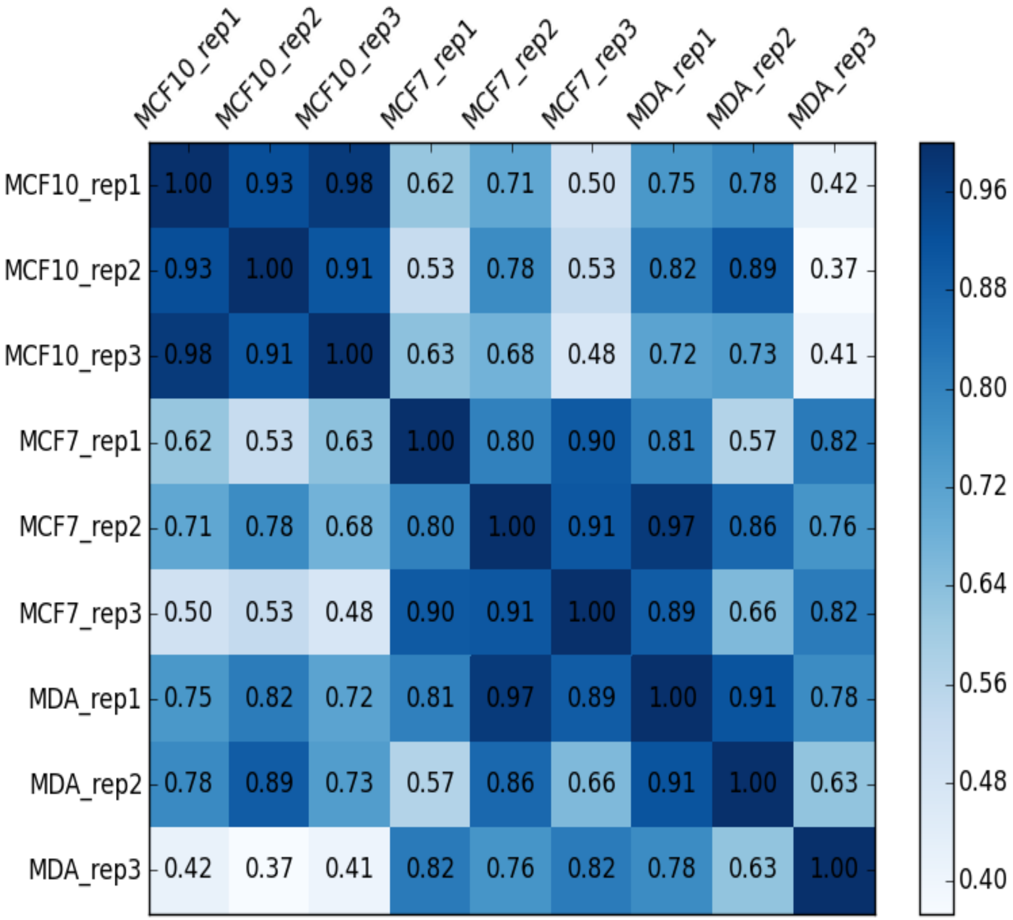


**Figure S1.** Spearman rank correlation between all 9 samples of the SF-miRNA collection identified in each sample.

**Table S1: Sequencing data for the breast cencer cell lines and replicates**

| Cell Replicates | Total reads | Mapped reads | % mapped reads | mapped to miRNA* | % of all mapped reads |
|---|---|---|---|---|---|
| MCF10A rep1 | 31150316 | 19173084 | 61.6 | 10611 | 0.055343 |
| MCF10A rep2 | 33718453 | 20087790 | 59.6 | 17053 | 0.084892 |
| MCF10A rep3 | 25493755 | 14013958 | 55 | 8611 | 0.061446 |
| MCF7 rep1 | 10716584 | 2242497 | 20.9 | 3687 | 0.164415 |
| MCF7 rep2 | 6492651 | 1725734 | 26.6 | 1882 | 0.109055 |
| MCF7 rep3 | 17231509 | 2600045 | 15.1 | 2957 | 0.113729 |
| MDA-231 rep1 | 11529138 | 2852117 | 24.7 | 4078 | 0.142982 |
| MDA-231 rep2 | 17987596 | 10005933 | 55.6 | 10571 | 0.105647 |
| MDA-231 rep3 | 11800122 | 2141920 | 18.2 | 6293 | 0.293802 |

* Reads of length >= 17 and at most 2 alignment to the genome

Supplementary Table S2, Table S3 and raw data for Table S4 are missing (too long for PDF)

**Table S4. Fraction of Mature of the other aligned reads by segmental region**

| | MCF10A | MCF7 | MDA-231 |
|---|---|---|---|
| extension | 13.89187 | 9.078925 | 8.727222 |
| mature | 25.44188 | 60.04101 | 68.23237 |
| undefined complement | 0.985612 | 1.026544 | 0.626379 |
| hairpin precursor | 3.027587 | 2.240941 | 1.245125 |
| overlap region | 56.65305 | 27.61258 | 21.1689 |

Removed extension that are beyond the complete complete hairpin premiRNA

**Table S5**. SF-miRNA expression in view of miRcancer

| Row.names | padj | Aver MCF-10A | Aver MCF-7 | Aver MDA-MB-231 | ratio MDA/MCF-10A | ratio MCF-10A/MDA | total MCF-10A+MDA | miRCancer evidence | Down (Cancer to healthy) | Up (Cancer to healthy) | our finding SF-miRNA | % Consistancy within miRCancer | Unified Annotation |
|---|---|---|---|---|---|---|---|---|---|---|---|---|---|
| hsa-mir-612 | 9.28E-34 | 450.45 | 1.57 | 3.05 | 0.007 | 147.920 | 453.50 | 2 | 2 | 0 | down | 1.000 | little support |
| hsa-mir-5047 | 6.26E-20 | 163.84 | 14.51 | 1.21 | 0.007 | 135.210 | 165.06 | 0 | 0 | 0 | down | NA | NA |
| hsa-mir-7161 | 1.51E-15 | 74.73 | 1.94 | 1.23 | 0.016 | 60.874 | 75.96 | 0 | 0 | 0 | down | NA | NA |
| hsa-mir-3687 | 1.39E-12 | 460.56 | 34.85 | 8.16 | 0.018 | 56.466 | 468.72 | 0 | 0 | 0 | down | NA | NA |
| hsa-mir-3064 | 4.10E-11 | 196.59 | 46.28 | 8.78 | 0.045 | 22.395 | 205.36 | 1 | 1 | 0 | down | 1.000 | little support |
| hsa-mir-1246 | 3.97E-10 | 484.63 | 83.84 | 32.00 | 0.066 | 15.144 | 516.63 | 10 | 3 | 7 | down | 0.3 | Mixed |
| hsa-mir-7704 | 5.07E-06 | 296.40 | 41.85 | 20.23 | 0.068 | 14.654 | 316.63 | 0 | 0 | 0 | down | NA | NA |
| hsa-mir-3607 | 6.75E-07 | 56.98 | 10.82 | 4.34 | 0.076 | 13.138 | 61.32 | 2 | 1 | 1 | down | 0.500 | little support |
| hsa-mir-1248 | 1.06E-07 | 66.78 | 4.54 | 5.25 | 0.079 | 12.719 | 72.03 | 0 | 0 | 0 | down | NA | NA |
| hsa-mir-148a | 1.71E-07 | 47.42 | 32.41 | 3.97 | 0.084 | 11.930 | 51.39 | 52 | 46 | 6 | down | 0.885 | Agree |
| hsa-mir-6087 | 4.74E-09 | 5847.12 | 1124.30 | 705.28 | 0.121 | 8.291 | 6552.40 | 0 | 0 | 0 | down | NA | NA |
| hsa-mir-1291 | 1.71E-07 | 88.03 | 147.59 | 14.91 | 0.169 | 5.903 | 102.95 | 3 | 2 | 1 | down | 0.667 | Mxed |
| hsa-mir-622 | 8.70E-05 | 54.58 | 74.17 | 16.81 | 0.308 | 3.246 | 71.40 | 9 | 7 | 2 | down | 0.778 | Agree |
| hsa-mir-21 | 1.06E-06 | 362.56 | 1173.27 | 1155.51 | 3.187 | 0.314 | 1518.07 | 268 | 7 | 261 | up | 0.026 | Agree |
| hsa-let-7f-1 | 1.67E-06 | 54.02 | 90.58 | 184.70 | 3.419 | 0.292 | 238.72 | 14 | 14 | 0 | up | 1 | Opposite |
| hsa-let-7f-2 | 7.02E-10 | 32.11 | 85.99 | 173.25 | 5.395 | 0.185 | 205.36 | 14 | 14 | 0 | up | 1.000 | Opposite |
| hsa-let-7i | 1.04E-06 | 21.19 | 62.31 | 140.25 | 6.618 | 0.151 | 161.45 | 17 | 17 | 0 | up | 1.000 | Opposite |
| hsa-mir-221 | 7.25E-14 | 8.52 | 0.00 | 59.69 | 7.007 | 0.143 | 68.21 | 98 | 9 | 89 | up | 0.092 | Agree |
| hsa-let-7g | 2.58E-05 | 20.37 | 84.80 | 170.57 | 8.372 | 0.119 | 190.94 | 20 | 19 | 1 | up | 0.950 | Opposite |
| hsa-mir-222 | 2.70E-26 | 6.46 | 0.00 | 105.45 | 16.328 | 0.061 | 111.91 | 61 | 6 | 55 | up | 0.098 | Agree |
| hsa-mir-30a | 2.73E-15 | 5.23 | 0.77 | 91.07 | 17.397 | 0.057 | 96.30 | 51 | 45 | 6 | up | 0.882 | Opposite |
| hsa-mir-100 | 4.65E-65 | 13.06 | 0.00 | 483.42 | 37.014 | 0.027 | 496.48 | 39 | 34 | 5 | up | 0.872 | Opposite |

At least 75% of evidence must be consistant. Otherwise marked as mixed evidence.

**Table S6. Expression trend for SF-miRNAs in breast cell lines and KM survival plots**

| SF-miRNA (22 genes) | miRNA MDA > MCF10[a] | HR | (95% confidence interval) | Logrank P-value | Data from KM Survival Plots[b] | Expected change from KM[c] | Inverse trend SF & KM plot |
|---|---|---|---|---|---|---|---|
| hsa-mir-6087 | D | 2 | 1.43-2.79 | 3.8e-05 | OncomiR | U | Opposite |
| hsa-mir-21 | U | 1.63 | 1.16-2.27 | 0.004 | OncomiR | U | Consistent |
| hsa-mir-1246 | D | 1.52 | 1.23-1.87 | 9e-05 | OncomiR | U | Opposite |
| hsa-mir-3687 | D | - | - | - | Not listed | - | |
| hsa-mir-100 | U | 0.7 | 0.57-0.85 | 0.00032 | Tumor sup. | D | Opposite |
| hsa-mir-612 | D | 2.06 | 1.48-2.87 | 1.5e-05 | OncomiR | U | Opposite |
| hsa-mir-7704 | D | 2.23 | 1.59-3.12 | 2.1e-06 | OncomiR | U | Opposite |
| hsa-let-7f-1 | U | 0.72 | 0.59-0.87 | 0.00079 | Tumor sup. | D | Opposite |
| hsa-let-7f-2 | U | 0.72 | 0.59-0.87 | 0.00079 | Tumor sup. | D | Opposite |
| hsa-let-7g | U | 0.72 | 0.59-0.88 | 0.0014 | Tumor sup. | D | Opposite |
| hsa-mir-3064 | D | 1.69 | 1.2-2.39 | 0.0025 | OncomiR | U | Opposite |
| hsa-mir-1291 | D | 1.35 | 0.96-1.9 | 0.088 | Not significant | n.s. | |
| hsa-let-7i | U | 0.82 | 0.66-1.01 | 0.065 | Not significant | n.s. | |
| hsa-mir-5047 | D | 2.23 | 1.59-3.12 | 2.1e-06 | OncomiR | U | Opposite |
| hsa-mir-622 | D | 2.23 | 1.59-3.13 | 2e-06 | OncomiR | U | Opposite |
| hsa-mir-3607 | D | 0.71 | 0.49-1.04 | 0.08 | Not significant | n.s. | |
| hsa-mir-1248 | D | 1.39 | 0.97-1.99 | 0.069 | Not significant | n.s. | |
| hsa-mir-222 | U | 1.35 | 1.07-1.71 | 0.013 | OncomiR | U | Consistent |
| hsa-mir-30a | U | 0.75 | 0.61-0.91 | 0.0041 | Tumor sup. | D | Opposite |
| hsa-mir-148a | D | 0.7 | 0.57-0.86 | 0.00062 | Tumor sup. | D | Consistent |
| hsa-mir-7161 | D | 2.2 | 1.57-3.06 | 3.3e-06 | OncomiR | U | Opposite |
| hsa-mir-221 | U | 1.24 | 0.87-1.76 | 0.23 | Not significant | n.s. | |

[a]Expression of SF-miRNA in MDA-MB-231 relative to MCF-10A cells. Trend is annotated U or D for up and down, respectively. [b]Data from KM survival plot for miRNAs from these cell lines. [c]Expected trend in these cell-lines as extracted from KM survival plots, where oncogenic miRNAs are annotated as U (up) while tumor suppressor miRNAs are expected to go down (D). miRNAs that do not exhibit a significant trend are marked as non-significant (n.s.). SF-miRNAs that result in inverse annotations for the cellular view and patients' survival are marked as opposite.
